# KLONE: An Imbalance-Robust *k*-NN Ensemble for Cross-Modal Single-Cell Clone Assignment^*^

**DOI:** 10.64898/2026.11.07.765116

**Authors:** Xiru Huang, Shengbin Ye, Arlei Silva, Luay Nakhleh

**Affiliations:** Department of Computer Science Rice University; Department of Statistics and Data Science Northwestern University

## Abstract

Single-cell DNA sequencing (scDNA-seq) and single-cell RNA sequencing (scRNA-seq) enable the inference of copy number clones and transcriptional states in tumor studies. However, these two assays usually profile different cells of the same tumor, requiring computational assignment of unpaired scRNA-seq cells to clones identified by scDNA-seq to link clonal genotypes with transcriptional profiles. This task is challenging because the only connection between the two modalities is a weak, gene-dependent dosage effect, and the absence of clone labels for RNA cells precludes supervised learning of this relationship. Moreover, clonal distributions may differ between the two modalities because the cells are profiled independently, while the inherent clonal imbalance of real tumors can leave minority subclones underrepresented, further compromising assignment accuracy. To address these challenges, we present KLONE, a per-clone *k*-nearest-neighbor (*k*-NN) ensemble that assigns RNA cells directly to copy-number clones based on cross-modal Pearson similarity, without the need for a parametric model or clone labels for RNA cells. To ensure a fair assessment of each clone’s compatibility with a given RNA cell, KLONE allocates an identical neighborhood budget to every clone, preventing majority clones from dominating the affinity assessment solely through their greater abundance. We compare KLONE with six existing methods on three colorectal cancer patients, using cells profiled in both modalities as ground truth. KLONE is the most accurate method and recalls minority clones at 100%. Despite discrepancies in clonal distribution between the two modalities, KLONE recovers the clonal distribution of RNA cells almost perfectly (*ρ* = 0.99).

## Introduction

Somatic copy number alterations (CNAs) are a hallmark of cancer [1]. A single CNA event often spans a chromosome arm or an entire chromosome, changing the copy number of hundreds of genes at once [2]. CNAs accumulate unevenly as a tumor grows, fracturing its cell population into genetically distinct subclones whose expansion and competition drive tumor evolution [3, 4, 5]. The clone is therefore the natural unit for studying a tumor, and single-cell sequencing resolves it directly. Single-cell DNA sequencing (scDNA-seq) reconstructs the clonal architecture from per-cell copy number profiles, now at a throughput of thousands of cells per tumor [6, 7, 8, 9]. Single-cell RNA sequencing (scRNA-seq), in turn, captures the transcriptional state of each cell, exposing the expression programs that distinguish tumor cell populations [10, 11].

Linking each subclone’s genome to its transcriptome would show how copy number alterations reshape transcriptional programs clone by clone [9, 12, 13], but genotype and phenotype are usually read out in *different cells*. Assays that sequence DNA and RNA from the same cell, such as G&T-seq [14], DR-seq [15], and scTrio-seq [16], profile far fewer cells than single-modality platforms [17], so large-scale tumor studies instead use high-throughput scDNA-seq and scRNA-seq platforms to sequence the two modalities from *separate cells* within the same tumor sample [8, 9, 12, 13]. Joint inference of such data thus requires *clone assignment*: given clones defined from the DNA cells, decide which clone each RNA cell belongs to. Three properties make this hard. First, the only link between these two modalities is gene dosage, and it is weak. Although a gene’s expression rises with its copy number on average [18], the relationship is less than proportional [19], differs from gene to gene [13, 20], and competes with regulatory and technical variation, so no single gene or cell pair is decisive. Second, because different modalities are measured in different cells, no RNA cell has a known clone, which makes supervised training and tuning impossible [21, 22]. Third, the two sets of cells are sampled independently, so the clonal distribution of the RNA cells need not be identical to that of the DNA cells [23, 24]. Compounding this challenge, tumors are often clonally imbalanced [6, 25], so a method that is not imbalance-robust tends to misassign the rare subclones to the dominant clone. This is particularly problematic because rare subclones can, for instance, drive treatment resistance [26, 27] or seed metastases [28], and correctly assigning RNA cells to them reveals the transcriptional programs behind these outcomes [27].

Existing methods differ in what they compare an RNA cell against, and each carries an assumption that does not hold for small, imbalanced tumor samples. *Clone-level dosage models* compare the cell against one consensus copy number profile per clone. clonealign [12] sets each gene’s expected expression proportional to its copy number in the clone, and TreeAlign [13] adds a per-gene switch that turns this dependence on or off. Both must estimate per-gene and per-cell parameters from the very cells they assign, and neither has a place for a gene that responds only partially. *Cell-level matching* compares the cell against individual DNA cells rather than against a consensus profile, so a clone is represented by the profiles of its member cells. Since both modalities sample the same tumor, the DNA cells include members of each RNA cell’s clone, and matching exploits this cross-modal correspondence by letting an RNA cell inherit the clone of the DNA cells it most resembles. MaCroDNA [24] does so without a parametric model. It showed that the Pearson correlation between an RNA cell’s expression and a DNA cell’s copy number already carries the clone signal, and it pairs RNA and DNA cells one-to-one to maximize the total correlation. The pairing, however, assumes that the two modalities share the same clonal distribution, and performance degrades when they do not [24]. *Population-level integrators* are general-purpose tools built to work across modality pairs by aligning the two sets of cells as wholes: Seurat [29] through mutual nearest neighbors in a shared projection, GLUE [30] through an adversarially aligned embedding, and MOSCOT [31] through an optimal transport (OT) coupling. Due to their general-purpose nature, they do not make use of the DNA-defined clone labels when aligning. Instead, they align the two datasets as if their compositions matched, and the OT coupling goes further by fixing the assigned clone proportions to the DNA panel’s, which degrades the alignment when the clonal distributions differ across modalities. Although GLUE corrects such differences with weights estimated from a preliminary clustering [30], our empirical results show that it is suboptimal among the tested integrators. No existing method overcomes all three challenges at once.

We introduce **KLONE**, a per-clone *k*-NN ensemble that addresses each of the three challenges above. It compares each RNA cell against individual DNA cells by Pearson correlation over their shared genes, so the weak dosage signal is pooled across many genes instead of being modeled by a noisy per-gene model. To fairly assess the fitness of each DNA clone to a given RNA cell, KLONE scores each clone by the *k* DNA cells of that clone most similar to the RNA cell. Every clone thus gets the same budget of *k* within-clone neighbors, and a larger clone gains no advantage from its size alone. Since no RNA labels exist for training or tuning, KLONE has no learned parameters and selects its configurations by an unsupervised entropy criterion. We evaluate KLONE and six existing methods on three colorectal cancer (CRC) patients [32], where some of the cells are profiled in both modalities and serve as ground truth for evaluation. On average, KLONE outperforms all six methods in both accuracy and macro-*F*_1_, and it assigns a patient in a fraction of a second. KLONE is also more robust to clonal imbalance than its competitors. In a controlled subsampling experiment that alters the clonal distribution, its minority clone recall remains at 100% as the majority clone grows to 92% of the cells. KLONE is open-source at https://github.com/NakhlehLab/KLONE.

## 2 Methods

### 2.1 The Clone Assignment Problem

Let 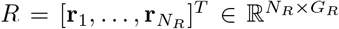 be the expression matrix and 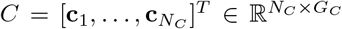 be the copy number matrix, such that 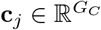 denotes the copy number of the *j*th DNA cell and 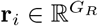 denotes the expression of the *i*th RNA cell. Each DNA cell carries a clone label *y*_*j*_ ∈ {1, …, *K*}, *K* ≪*N*_*C*_, inferred upstream by clustering the DNA cells’ copy number profiles and taken as given. The corresponding clonal distribution of the DNA panel is thus 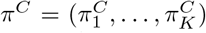, where 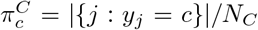 is the proportion of clone *c* in the DNA panel. Given (*R, C*, **y**), the clone assignment problem aims to infer the latent clone labels 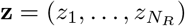 of the RNA cells. Since *K* ≪ *N*_*R*_, many RNA cells share a clone and the assignment is thus many-to-one. Furthermore, *N*_*R*_ and *N*_*C*_ are not necessarily equal, which precludes any rule that pairs each RNA cell with a distinct DNA cell, as it would be structurally misspecified.

The three difficulties from Section 1 take a precise form in this notation. First, the cross-modal coupling is weak. Copy number exerts an attenuated positive influence on expression [18, 19, 20], so we assume only that cells of the same clone are more cross-modally correlated than cells of different clones once evidence is aggregated over many genes. That is, E[*ρ*_*ij*_ |*z*_*i*_ = *y*_*j*_] *>* E[*ρ*_*ij*_ |*z*_*i*_ ≠ *y*_*j*_], where *ρ*_*ij*_ is the cross-modal Pearson similarity of Eq. (1). The margin is thin, as the per-patient median gap between same- and different-clone correlations is only 0.006 to 0.09, so no single cell pair is reliable. Second, RNA clone labels **z** are not observed, so there is nothing to train on and no validation set to tune against. Third, since the RNA and DNA cells are sampled independently, the RNA clonal distribution 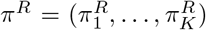, where 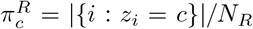, need not match the DNA clonal distribution *π*^*C*^.

Thus, a *frequency-constrained* rule, one whose assignments depend on the observed *π*^*C*^, is misspecified and leads to incorrect assignments. Instead, the inferred RNA clonal distribution 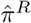 should track the unobserved *π*^*R*^.

KLONE answers each difficulty with one design choice. It meets the weak coupling by pooling the dosage signal across many genes into a single Pearson similarity, and the many-to-one structure by replacing matching with a soft per-cell vote (Section 2.2). It removes the dependence on *π*^*C*^ by scoring each clone on its own nearest neighbors with an equal per-clone budget, so the larger clones do not dominate the neighborhood and bias the assignments (Section 2.3).

Furthermore, it bypasses the need for the latent RNA clone labels **z** by fixing a grid of configurations and selecting among them with an unsupervised entropy criterion (Sections 2.4–2.5). Figure 1 overviews the KLONE pipeline.

**Figure 1:**
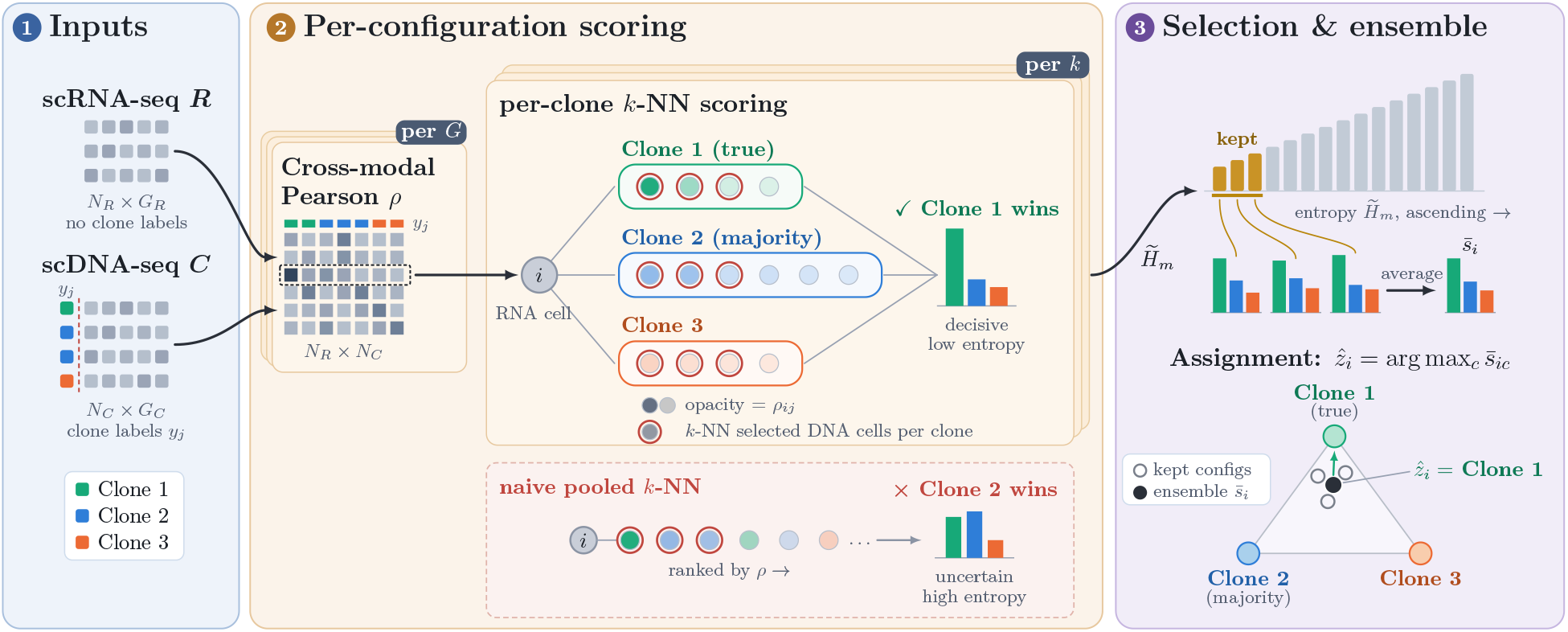
Overview of KLONE on a three-clone example in which RNA cell *i* belongs to the minority clone 1. Green, blue and orange denote clones 1 to 3 throughout. **(1)** Inputs. An scRNA-seq count matrix ***R*** (*N*_*R*_ cells, no clone labels) and an scDNA-seq copy-number matrix ***C*** (*N*_*C*_ cells) with clone labels *y*_*j*_ from clustering the copy-number profiles. **(2)** Per-configuration scoring. For each gene budget *G* and neighbor count *k* on a fixed grid, KLONE computes the cross-modal Pearson similarity *ρ*_*ij*_ over the top-*G* highly variable genes (Eq. 1) and scores each clone by the mean positive similarity of cell *i*’s *k* nearest DNA cells within that clone (Eq. 3): opacity denotes affinity and red rings mark the *k* picks. Since every clone receives the same budget, one strong clone-1 neighbor outweighs several medium clone-2 neighbors and the score is decisive, whereas a naive pooled *k*-NN over all DNA cells (bottom box) lets two medium clone-2 neighbors outvote it, so the majority clone wins with an uncertain, high-entropy score. **(3)** Selection and ensemble. KLONE ranks the configurations by standardized mean entropy 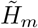 (Eq. 4), keeps the *N*_pc_ lowest (gold), averages their score vectors into 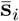 and assigns 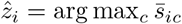 (Eq. 5). On the simplex, open circles are the kept configurations and the filled dot is their average, nearest the clone-1 vertex. No step uses RNA clone labels.

### 2.2 Cross-Modal Similarity and Soft *k*-NN Voting

Motivated by the many-to-one structure of clone assignment (Section 2.1), KLONE replaces explicit correspondences with a soft, *local k*-NN vote. We measure RNA–DNA cell similarity by cross-modal Pearson correlation, which captures the gene-dosage relationship. Furthermore, it is insensitive to per-cell shifts and scales that differ across the two modalities. Specifically, for a gene set *G* shared among both modalities, the similarity between RNA cell *i* and DNA cell *j* is measured by

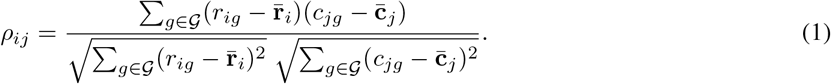

Given the RNA–DNA cell similarity matrix 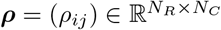, each RNA cell *i*’s clone assignment depends on its *k* nearest DNA cells 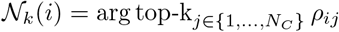. We define *s*_*ic*_, the normalized affinity of RNA cell *i* to clone *c*, as the total positive similarity coming from its neighbors in clone *c*:

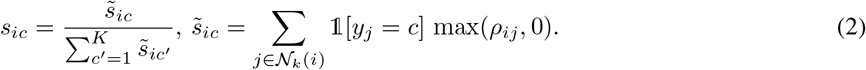

In the degenerate case where 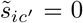 for all 1≤ *c*^*′*^≤ *K*, we define *s*_*ic*_ = 0. By construction, **s**_*i*_ = (*s*_*i*1_, …, *s*_*iK*_) is a probability vector on the *K*-simplex Δ^*K*−1^, where only positive similarities vote, so anti-correlated neighbors contribute nothing. Unlike a hard match, this vote is local to each RNA cell, aggregates evidence over *k* neighbors, and returns a soft score *s*_*ic*_. That is, the same DNA cell can be assigned to many RNA cells, if the evidence supports it, and the assignment of one RNA cell does not affect the available DNA cells for the other.

### 2.3 Per-Clone *k*-NN Scoring

The pooled vote of Section 2.2 draws *N* _*k*_(*i*) from *all* DNA cells. Under a heavily imbalanced panel, the neighborhood *N* _*k*_(*i*) can be infiltrated by the majority clone for most RNA cells, i.e., the frequency-constrained bias of Section 2.1, which skews the assignments.

To circumvent this clonal imbalance bias issue, we introduce the per-clone *k*-NN score 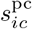, which removes this bias by giving every clone the same neighbor budget. Let *C*_*c*_ = {*j* : *y*_*j*_ = *c*} denote the set of DNA cells belonging to clone *c, n*_*c*_ = |*C*_*c*_| be the number of DNA cells in clone *c*, and 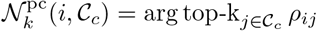 be the *k* nearest clone-*c* DNA cells of RNA cell *i*. We define the clone-*c k* -NN score of RNA cell *i* as

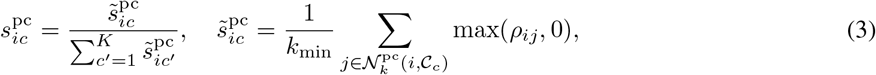

where *k*_min_ = min(*k, n*_*c*_). That is, we restrict the *k*-NN search to *C*_*c*_, average the positive similarity *within* clone *c*, and normalize across all clones to produce a probability vector 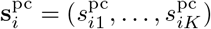 on the *K*-simplex Δ^*K*−1^.

By construction, 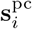 is insensitive to the clone size *n*_*c*_ since each clone’s neighbors are drawn independently, so a minority clone competes on the quality of its own best neighbors rather than being outvoted by a dominant one. The component ablation in Section 3.6 confirms that 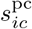 helps recover minority-clone performance under imbalance. Furthermore, since 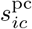, is independent of the clone size *n*_*c*_ the inferred RNA clone-*c* proportion 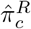 is not constrained to match the observed DNA clone-*c* proportion 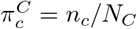. In contrast, MOSCOT fixes the transported marginal to *π*^*C*^ and MaCroDNA caps the number of RNA cells assigned to clone *c* at *n*_*c*_, so both reproduce *π*^*C*^ rather than *π*^*R*^ (Supp. K). Section 3.5 shows that KLONE’s 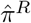 tracks *π*^*R*^ even when the majority clone reaches 91%.

### 2.4 Entropy-Based Configuration Selection

Both the neighborhood size *k* and the gene budget *G* = |*G*| (the number of highly variable genes used to compute the similarity; see Supp. A) are hyperparameters that trade bias against variance and signal against noise, and the best choice is patient-dependent (Section 3.1). Crucially, we cannot pick *G* or *k* by validation, because the RNA cells are unlabeled.

To this end, we design a label-free, entropy-guided configuration selection module that sweeps a grid of (*G, k*) configurations, which removes (*G, k*) as hyperparameters instead of tuning them. Specifically, we evaluate a fixed grid of *M* configurations over gene budgets *G* and neighbor counts *k*, each producing per-clone score vectors 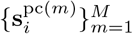.

We measure how decisively each configuration commits by the mean Shannon entropy of its per-cell scores, computed over all *N*_*R*_ unlabeled RNA cells,

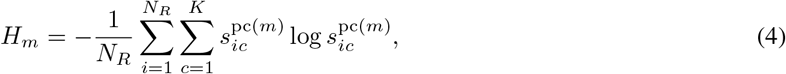

where the degenerate case 0 log 0 0. We choose Shannon entropy because it is the canonical label-free measure of how decisively a score vector commits: for an RNA cell *i*, it is minimized when the mass concentrates on one clone and maximized when spread uniformly over the *K* clones. A low *H*_*m*_ therefore marks a configuration that resolves the cross-modal similarity into sharper assignments than its peers.

Nonetheless, the raw entropy *H*_*m*_ is not directly suitable here. Since *k* sets how many neighbors are averaged into each clone score, it determines what the score vector *is* and not merely how decisive it is. Thus, a configuration’s entropy *H*_*m*_ is informative relative to those built with the same *k*. Let *μ*_*k*_ and *σ*_*k*_ denote the mean and standard deviation of {*H*_*m*_ : *k*_*m*_ = *k*}. We rank configurations by the standardized entropy 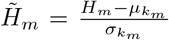 and retain the lowest *N*_pc_ standardized entropy 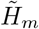 for model ensembling.

Two properties follow. First, selection is fully unsupervised: 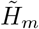 depends only on the geometry of the unlabeled scores and never requires the unknown RNA labels. Second, since 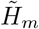 is recomputed per patient, KLONE adopts the gene budget *G* on which each patient’s dosage signal is sharpest rather than committing to a single global default.

### 2.5 The KLONE Ensemble

Let *T*_pc_ index the *N*_pc_ = 3 most decisive configurations. The final assignment averages the *T*_pc_-selected per-clone score vectors and takes the most likely clone:

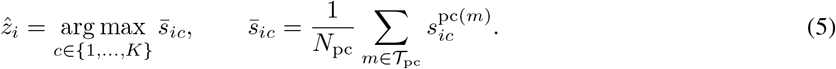

Since each 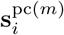 lives on the simplex Δ^*K*−1^, the average 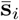 is a convex combination that remains a valid clone-membership distribution. In particular, the predicted label 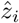 is the mode. Supp. Algorithm 1 summarizes the KLONE pipeline.

Averaging the selected 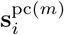 reduces variance. A single configuration’s score is noisy: a large *G* can admit a buffered amplicon that pulls it off-target, and a small *k* can let one outlying neighbor dominate. Because the retained configurations use different (*G, k*), their errors rarely align, so averaging cancels configuration-specific noise while preserving the clone signal they share. We average only the *N*_pc_ lowest-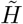 configurations because the discarded higher-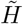 ones have diffuse scores that would blur the decisive assignments. Furthermore, KLONE is robust to the choice of *N*_pc_ (Supp. H).

KLONE is also closed form. For one configuration, the similarity matrix in Eq. (1) costs *O*(*N*_*R*_*N*_*C*_*G*) and the per-clone scores in Eq. (3) an additional *O*(*N*_*R*_*N*_*C*_) for the within-clone top-*k* selection. In total, evaluating *M* configurations costs *O*(*MN*_*R*_*N*_*C*_*G*_max_), with *G*_max_ the largest gene budget in the grid. Furthermore, no step in KLONE requires iterative parameter fitting, so assigning a patient only takes a fraction of a second (Section 3.7).

## 3 Results

### 3.1 Experimental Setup

To assess KLONE’s performance against other methods, we apply them to the colorectal cancer (CRC) cohort of [32], generated with single-cell triple-omics sequencing (scTrio-seq2), which jointly measures copy number and gene expression. We use three patients, `crc04`, `crc10`, and `crc11`. The rest are excluded for a differing library protocol or too few dual-modality cells (Supp. A). We restrict both modalities to the 12,894 genes measured in both modalities.

We evaluate on *anchor cells*: a subset of cells with identical barcodes in both modalities. Because their RNA and CNA profiles come from the same physical cell, we attach the clone label *y*_*i*_ to the latent RNA label *z*_*i*_, giving a ground-truth evaluation set under real biological conditions rather than a synthetic benchmark whose realism is hard to guarantee [21, 33]. Anchor RNA cells are among those being assigned, but their clone labels are held out from every method and used only for evaluation.

The clones themselves are not directly observed: a tumor’s clones are latent populations inferred by clustering the scDNA-seq copy number profiles, so the assignment targets depend on an uncertain upstream choice of clustering algorithm and data-preprocessing steps. That is, a method that looks accurate under one clustering may merely be exploiting that partition. To ensure our evaluation is not biased toward a single partition, we assess all methods across four clone definitions by crossing two copy number transformations (raw and log) with two clustering strategies: intNMF [34], which selects *K* automatically (*K* = 2 or 3 in our datasets), and agglomerative clustering fixed at *K* = 4 to capture finer subclonal structure [25, 35]. Reporting across all four definitions measures not just accuracy but *robustness* to this upstream clonal inference strategy.

We compare against six methods plus a trivial floor, ranging from clone assignment methods to general cross-modal integrators. Among the clone assignment methods, **MaCroDNA** performs *one-to-one matching*, solving a parameter-free ILP on Pearson correlation over all common genes, while **clonealign** [12] and **TreeAlign** [13] are probabilistic models in which a gene’s expression depends on its copy number in the cell’s clone. The general cross-modal integrators utilize different techniques: **Seurat-CCA** represents the *anchor-based* tools, building canonical correlation analysis (CCA) anchors to transfer labels, **GLUE** represents *deep generative* integration (graph-linked VAE), and **MOSCOT** represents *OT*. TreeAlign and clonealign may leave a cell unassigned, so we evaluate each cell’s maximum a posteriori (MAP) clone and report the packages’ default thresholded outputs in Supp. E. **Trivial** assigns every RNA cell to the majority clone, the floor any useful method must clear. Each baseline runs at its recommended settings. Exact preprocessing and configurations are deferred to Supp. A and D.

We report two per-cell metrics on the anchor cells, *accuracy* and *macro-F*_1_. Among the ground-truth anchor cells, the majority clone ranges from a mildly skewed 57.9% in `crc04` to an extremely imbalanced 91.4% in `crc11` (see Supp. Table S1). Thus, accuracy alone is misleading: a method that merely follows the dominant clone already scores 91.4% on `crc11`. Macro-*F*_1_ instead weights every clone equally and exposes minority-clone performance. We also measure *clonal-distribution recovery*, the Pearson correlation between *π*^*R*^ and 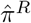 computed over the anchor cells, a distinct axis on which a method can misplace individual cells yet still reproduce the proportions, or vice versa.

#### KLONE configuration

KLONE sweeps a fixed grid of *M* = 15 configurations, crossing gene budgets *G* ∈ {500, 750, 1000, 1500, 2000} with neighbor counts *k* ∈ {2, 3, 5}, and averages the *N*_pc_ = 3 most decisive configurations. None of them is tuned. KLONE remains the best performer when varying *N*_pc_ ∈{2, 3, 5, 8, 10, 13} (Supp. H). The same holds when comparing KLONE to the baselines at their optimal *G* (Supp. I).

### 3.2 KLONE is the Most Accurate Method in Most Cases

KLONE assigns RNA cells to their correct clone more accurately than every baseline, on most patients (Figure 2, left). Averaged over the four clone definitions and three patients, it reaches 92.5% accuracy, 7.0 points above the next-best method, MaCroDNA (85.5%), and well above Seurat-CCA (81.5%), MOSCOT (78.5%), GLUE (75.9%), and the trivial floor (65.8%), which neither TreeAlign (65.4%) nor clonealign (38.7%) clears, as both misassign all RNA cells to a single clone in some runs (Supp. F). Both models estimate their per-gene dosage effects from the very cells they assign, so the fit admits degenerate optima in which one clone absorbs every cell. clonealign is the worse of the two, collapsing in eight of the twelve runs. Because expression responds less than proportionally to copy number [36], its proportional dosage model favors the clone with the flattest copy number profile, in four runs one that holds no anchor cells at all (its 0% runs). TreeAlign does not clear the trivial floor only on `crc04` (42.1% vs 57.9%), where the clonal distributions of the DNA and the RNA cells differ, funneling every RNA cell into the DNA panel’s largest clone, the anchor cells’ minority.

**Figure 2:**
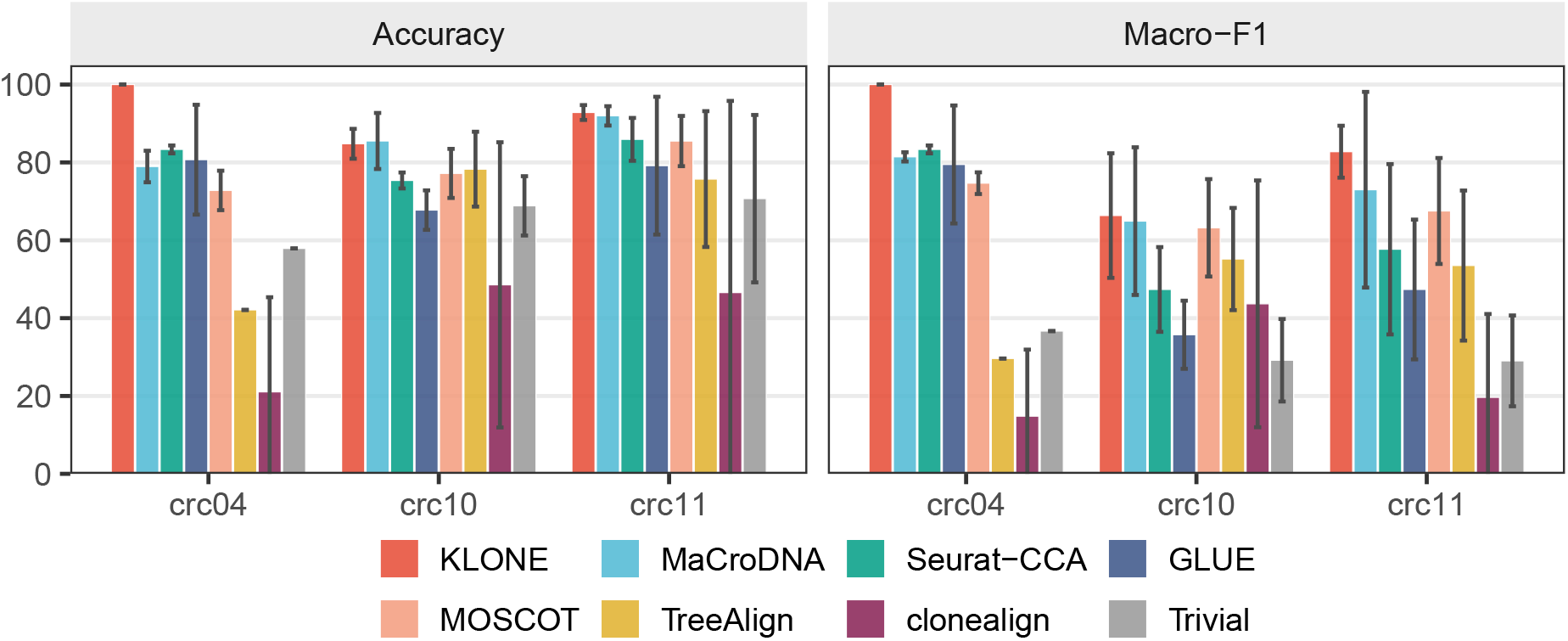
Per-patient accuracy (left) and macro-*F*_1_ (right) on the anchor cells of `crc04`, `crc10` and `crc11` for KLONE, the six baselines and the Trivial majority-clone reference. Bars are the mean over the four clone definitions and whiskers span *±*1 SD across them. TreeAlign and clonealign are scored on their MAP clone (Supp. E).

Among the three patients, the margin is widest on `crc04`, where KLONE reaches 100% accuracy under every clone definition while MaCroDNA falls to 78.9% as its one-to-one matching is forced to spread a few cells across distinct DNA cells and inevitably misroutes some. On the more imbalanced `crc10` and `crc11`, KLONE and MaCroDNA converge on accuracy (84.8% vs 85.5% and 92.8% vs 92.0%), while the remaining methods trail far behind (48.6–78.3% on `crc10` and 46.6–85.9% on `crc11`). KLONE is also the steadiest. Its per-patient accuracy barely moves as the clone definition changes (average SD 1.9 vs 2.9–36.7 for the other methods), so its lead does not hinge on any one way of defining the clones.

Accuracy alone, however, flatters the other methods on the imbalanced patients (Figure 2, right). On `crc11`, Seurat-CCA, MOSCOT, and GLUE achieve 79.2–85.9% accuracy yet only 47.4–67.5% macro-*F*_1_, the signature of following the majority clone. On the other hand, KLONE holds 82.7% in macro-*F*_1_ on `crc11`, the strongest among all competing methods. The same story holds when averaging across the cohort: KLONE’s macro-*F*_1_ (83.0%) leads all other methods, ahead of MaCroDNA (73.1%) by 9.9 points, MOSCOT (68.5%) by 14.5, and Seurat-CCA (62.8%), GLUE (54.2%), TreeAlign (46.1%), and clonealign (26.0%) by more than 20. The trivial floor itself is weaker on macro-*F*_1_ (31.6%), as the majority vote scores 0 *F*_1_ on every minority clone. TreeAlign clears it (46.1%) through the minority-clone cells it recovers on `crc10` and `crc11`, while clonealign (26.0%) stays below it on every patient except `crc10` (Supp. F).

These gaps are statistically significant. Bootstrap 95% CIs on the grand-mean metrics separate KLONE from every baseline, including the closest, MaCroDNA (macro-*F*_1_ [78.2, 84.8] vs [70.1, 75.3], accuracy [90.9, 94.0] vs [83.2, 87.7]).

A per-cell McNemar test over all anchor-cell assignments rejects equality with every baseline at Holm-adjusted *p <* 10^−4^. A conservative run-level Wilcoxon test over the 12 patient × clustering evaluations agrees, ranking KLONE above MaCroDNA (adjusted *p* = 0.046 for macro-*F*_1_, 0.045 for accuracy) and above every other baseline at adjusted *p <* 0.005 on both metrics (Supp. J).

### 3.3 Robust to the Clone Definition

Because the true clones are unknown in advance, a method cannot choose the clustering it will run under, so its worst-case behavior is what matters in practice. KLONE is both the most robust to the clone definition and often the most accurate under any single one (Figure 3). Its weakest definition still reaches 91.7% average accuracy, above the *best* definition of every baseline (MaCroDNA 87.4%, Seurat-CCA 82.9%, GLUE 83.8%, MOSCOT 81.2%, TreeAlign 72.2%, clonealign 64.7%), and it stays in a tight 91.7–93.4% band while the learned baselines are brittle (e.g., GLUE swings 12.6 points across definitions). The gap widens on macro-*F*_1_. Every method except clonealign, which scores 0% under agglomerative-log, scores lowest under agglomerative-raw, where KLONE still holds a 75.2% floor, 15.3 points above MaCroDNA’s (59.9%) and above the best macro-*F*_1_ that Seurat-CCA (70.4%), GLUE (62.9%), TreeAlign (51.8%), or clonealign (45.4%) reach under *any* definition. With no parameters or labels to fit, KLONE has nothing for a particular partition to destabilize, as the short whiskers in Figure 2 already foreshadow, whereas the model-based baselines must re-fit to each clustering.

**Figure 3:**
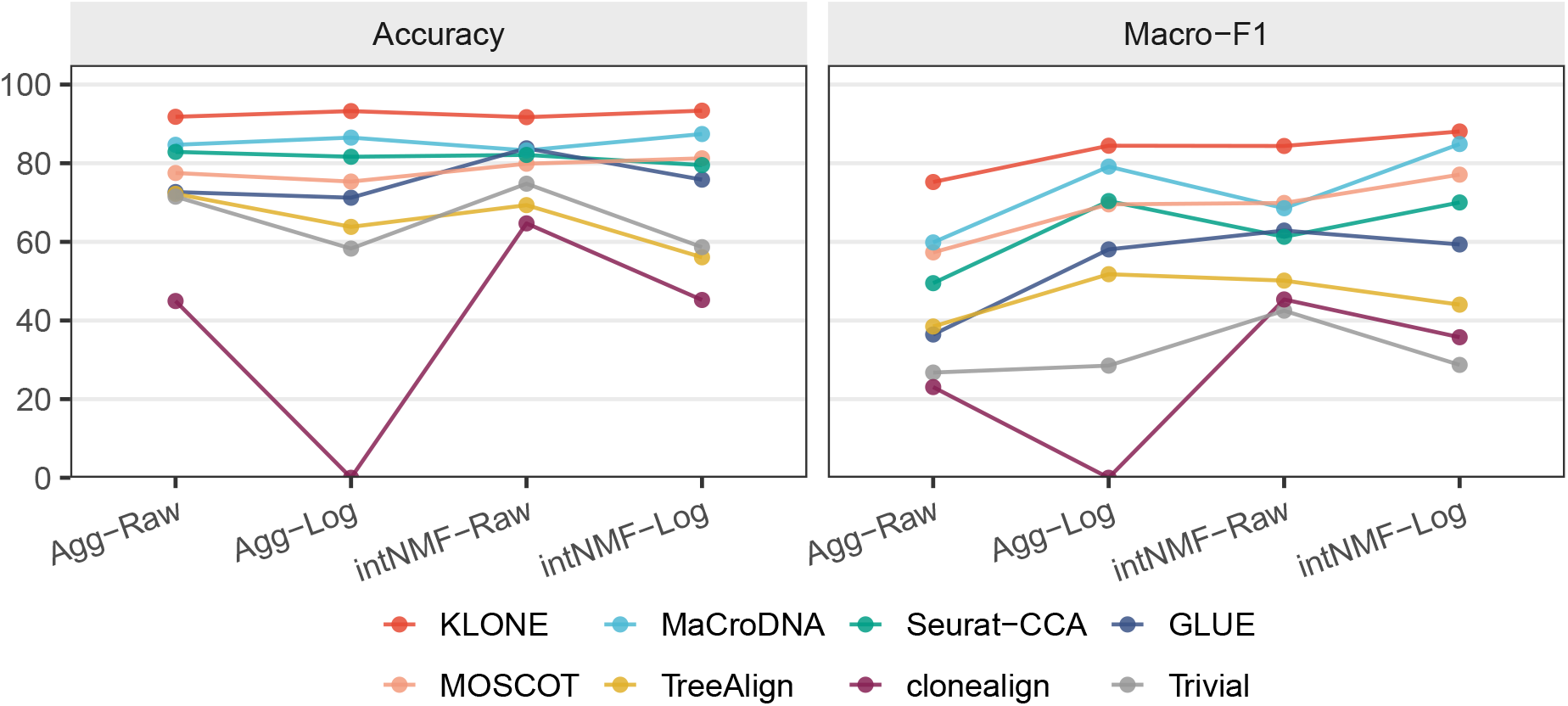
Accuracy (left) and macro-*F*_1_ (right) of the eight methods under each of the four clone definitions, agglomerative (Agg) or intNMF clustering on raw or log copy number, one line per method. Points are the per-patient score on anchor cells averaged over the three patients.

### 3.4 Robust to Clonal Imbalance

As shown in Figure 2, KLONE is the best performer even in clonal imbalance regimes (`crc10` and `crc11`). To validate this claim rigorously, we run a controlled experiment on `crc04` by randomly subsampling its minority clone cells, allowing us to vary the majority clone proportion *π*_max_ while retaining real biology signals (Supp. K). We use `crc04` since it is the least imbalanced patient, allowing us to simulate from a near-balanced baseline to extreme imbalance. To keep the minority clone from collapsing, we cap *π*_max_ at 0.92 so there are at least 3 minority clone anchor cells to evaluate on. We run each method over 30 resampled replicates per *π*_max_ and report the average macro-*F*_1_ and minority clone recall along with their bootstrap 95% CI. Empirically, KLONE achieves 100% macro-*F*_1_ in 190*/*210 instances (Figure 4), while the remaining 20 all score 99.23%, each misassigning one majority clone cell into a distractor clone. Similar observations hold for the minority clone recall, so the correct clone supplies the winning neighbors for nearly all RNA cells, no matter how rare that clone becomes. By contrast, the baselines behave differently and `crc04` is not an easy case for them. Though their failure modes differ slightly (Supp. K), the macro-*F*_1_ of most baselines drops as *π*_max_ increases, primarily due to a worsening minority clone recall as the minority clone becomes rarer (Figure 4). The exceptions, TreeAlign and clonealign, collapse instead. TreeAlign assigns every anchor cell to the minority clone up to *π*_max_ = 0.75. From *π*_max_ = 0.80, the minority clone has fewer than 10 DNA cells, so TreeAlign’s own clone-size filter drops it, and its macro-*F*_1_ rises while its minority clone recall falls to 0. clonealign sends every cell to a flat, diploid distractor clone in each subsample it fits, so its macro-*F*_1_ is 0 at every *π*_max_.

**Figure 4:**
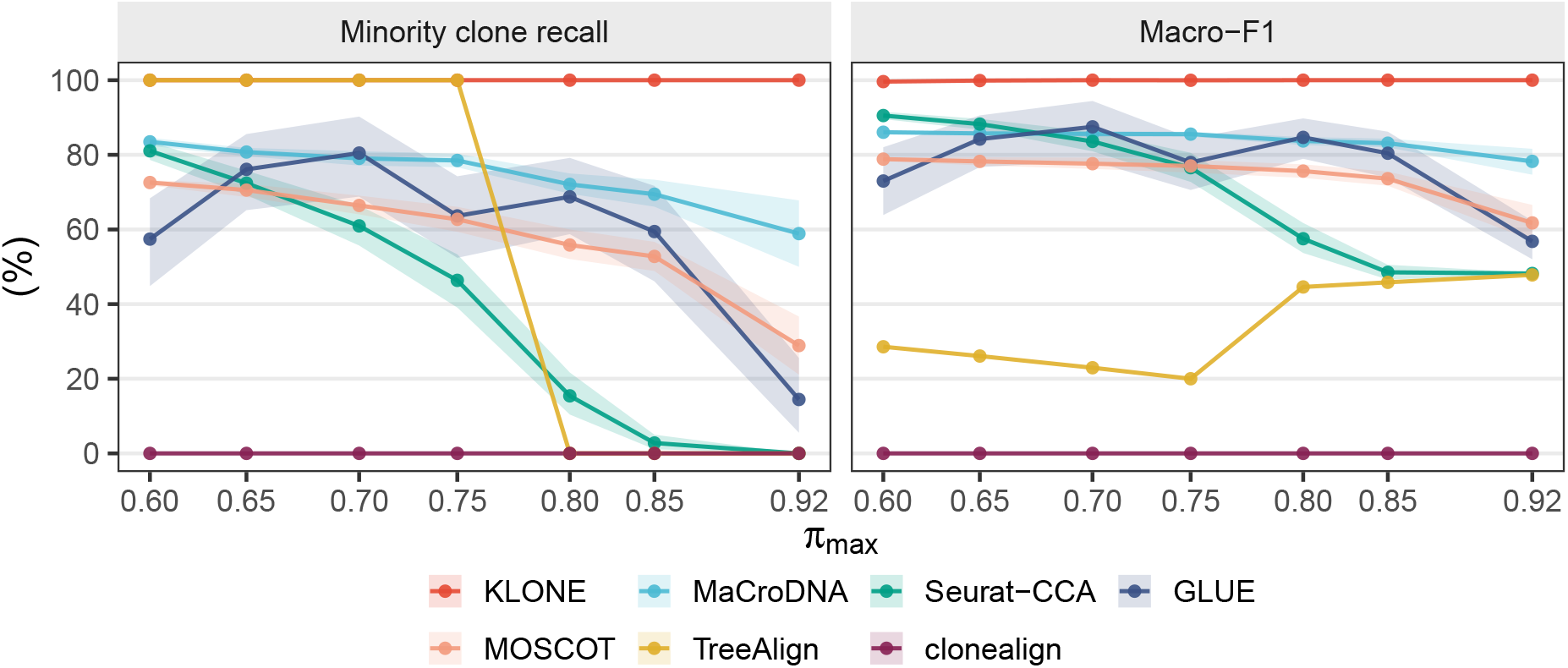
Imbalance-robustness sweep on `crc04` under the agglomerative-raw clone definition. Minority clone recall (left) and macro-*F*_1_ (right) of the seven methods versus the majority clone proportion *π*_max_. Points are the mean over 30 replicate subsamples per *π*_max_ and ribbons their bootstrap 95% CI. Minority clone size varies from 22 to 3 anchor cells.

### 3.5 KLONE Recovers the Clonal Distribution

Recovering the clonal distribution is a distinct axis from per-cell accuracy since a method can excel at one while missing the other. To measure how well a method recovers the clonal distribution, we compute the Pearson correlation between *π*^*R*^ and 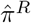 across clones and patients (Supp. L).

KLONE recovers the clonal distribution almost perfectly, and more faithfully than every baseline (Figure 5A). Its mean correlation across the four clone definitions is 0.99 vs 0.95 for the next best method, Seurat-CCA. This reflects KLONE tracking *π*^*R*^ rather than *π*^*C*^: its 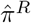 sits within 3.5% total-variation distance of *π*^*R*^, whereas the frequency-constrained MaCroDNA and MOSCOT land closer to *π*^*C*^ than to *π*^*R*^ (Supp. L). KLONE is also the most stable in this task. Across the four clone definitions, its correlations stay between 0.992 and 0.997, a spread of 0.005 versus 0.041 for the next most stable baseline, Seurat-CCA, whereas the other methods scatter widely across definitions.

**Figure 5:**
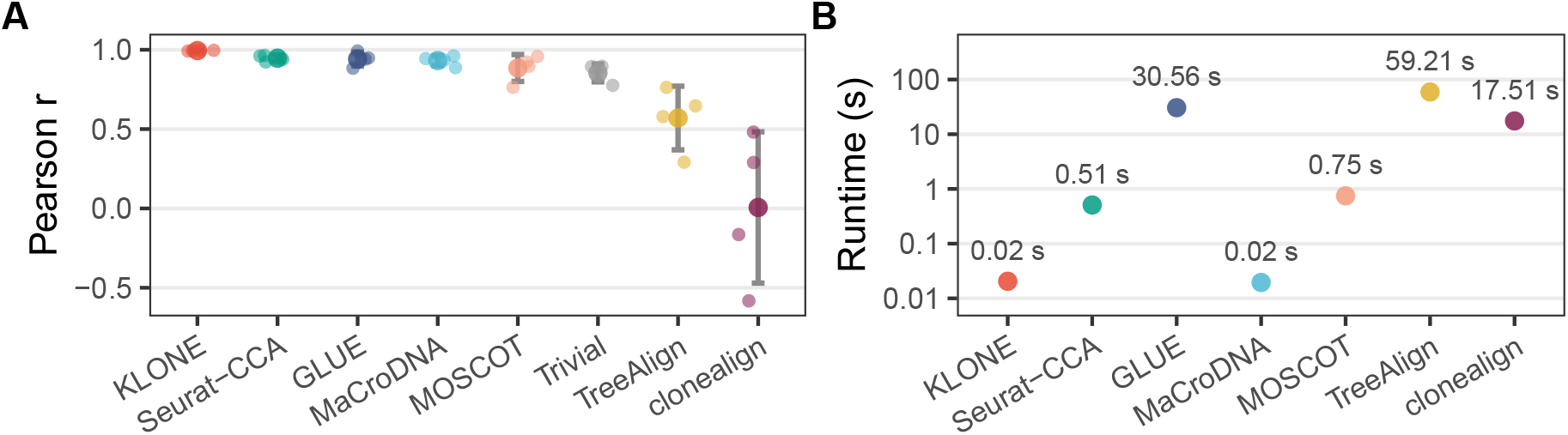
Clonal-distribution recovery and runtime of the eight methods, ordered by mean Pearson correlation. Pearson correlation between *π*^*R*^ and 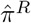 across clones and patients. Small dots represent the Pearson correlation per clone definition, and large dots represent the overall mean. Whiskers span *±* 1 SD. **(B)** Mean inference runtime per patient in seconds (log scale) over the 12 clustering × patient runs, on an Apple M4 Max (16-core) CPU with 8 threads for every method.

This is the clonal-distribution signature of per-clone scoring: a method that collapses onto the majority clone can still score decently on accuracy yet necessarily miscalibrate the clonal distribution. By contrast, scoring each clone on its own neighbors keeps the minority abundances intact, which is essential for downstream analyses that focus on rare subclones, such as therapy resistance and metastasis [4, 5].

### 3.6 Ensembling and Per-Clone Scoring Each Contribute

To isolate KLONE’s two design choices, per-clone scoring and ensembling of entropy-selected configurations, we compare it against three strategies that remove one or both while holding everything else fixed (Table 1). The *standardvote ensemble* drops the per-clone scoring with the pooled *k*-NN vote of Eq. (2), whose neighbors are drawn from *all* DNA cells at once. The *per-clone single configuration* drops ensembling, keeping the per-clone vote but only the single most decisive configuration (*N*_pc_ = 1). The *standard-vote single configuration* removes both.

**Table 1:** Component ablation. Values are grand means over the four clone definitions (%). The *agg-raw* column represents the most imbalanced definition, where majority bias is most punishing. Best in **bold**.

|  |  | Macro-*F*<sub>1</sub> |  |
|---|---|---|---|
| Strategy | Accuracy | mean | agg-raw |
| Standard-vote, single config | 91.3 | 81.3 | 72.6 |
| Standard-vote, ensemble | 90.9 | 78.1 | 64.0 |
| Per-clone, single config | 91.7 | 81.9 | 74.5 |
| **KLONE** | **92.5** | **83.0** | **75.2** |

Per-clone scoring is the dominant choice, and the two interact. At a single configuration, it lifts macro-*F*_1_ only modestly over the standard vote (81.3% to 81.9%). Its value appears under ensembling: because the pooled vote is swamped by the majority clone, averaging its three lowest-entropy configurations lowers macro-*F*_1_ from 81.3% to 78.1%, whereas averaging the de-biased per-clone scores (KLONE) raises it from 81.9% to 83.0%. The two choices are therefore complementary, and ensembling pays off only after per-clone scoring removes the majority bias. On the most imbalanced definition (agglomerative-raw) the split is starkest, the standard-vote ensemble collapsing to 64.0% macro-*F*_1_ while KLONE holds 75.2%. Crucially, the label-free KLONE loses almost nothing to supervision: it reaches 92.5% accuracy, converging to a per-patient oracle that picks each patient’s single accuracy-optimal configuration using the held-out labels (92.8%), and its 83.0% macro-*F*_1_ falls within 1.6 points of that oracle’s (84.6%). Averaging the three most decisive configurations is what closes this gap: the single most decisive pick reaches only 91.7%, so the ensemble, not any single label-free choice, closes the gap to the oracle (Supp. G).

### 3.7 Computational Efficiency

We evaluate the runtime of every method on an Apple M4 Max (16-core) CPU, where each method is given 8 threads. Figure 5B reports the average runtimes for the empirical experiment outlined in Section 3.1, where each method is averaged over the 12 clustering ×patient runs. KLONE is among the fastest methods, taking 0.02 s to assign a patient, on par with MaCroDNA (0.02 s) but 26 ×faster than the next fastest baseline, Seurat-CCA (0.51 s), and 2,961 ×faster than the slowest method tested, TreeAlign (59.21 s). The gap is structural: both KLONE and MaCroDNA are closed-form and require no model fitting, whereas the other baselines pay for their modeling machinery, and yet, gain no accuracy.

The sub-second runtime of KLONE is also what makes the clonal imbalance experiment of Section 3.4 cheap. Re-running KLONE under all four clone definitions and the 210-subsample imbalance sweep costs 2.58 s in total, whereas the least accurate and the third slowest method, clonealign, alone took 72 minutes. Moreover, KLONE’s speed enables it to be paired with any upstream clone inference method rather than committing to one.

### 3.8 Divergent Genotypes Yield Convergent Transcriptomes

Because KLONE places each RNA cell on a specific clone, we can ask how the clones’ genomic differences propagate into their transcriptomes. `crc10`’s clones are defined largely by a focal amplification of the HER2/ERBB2 oncogene (17q12): the amplicon rises from two copies in the diploid clone 1 to 20–33 copies in the others, spanning a 2-to-38-copy range across cells. Yet, almost none of this dosage reaches the transcriptome (Figure 6C). The expression climbs only 0.019 log-CPM per copy doubling, 6.3× weaker than for a dosage-concordant chromosome-12 control (0.119). This buffering renders the feature that most cleanly separates `crc10`’s clones in DNA nearly invisible in RNA.

**Figure 6:**
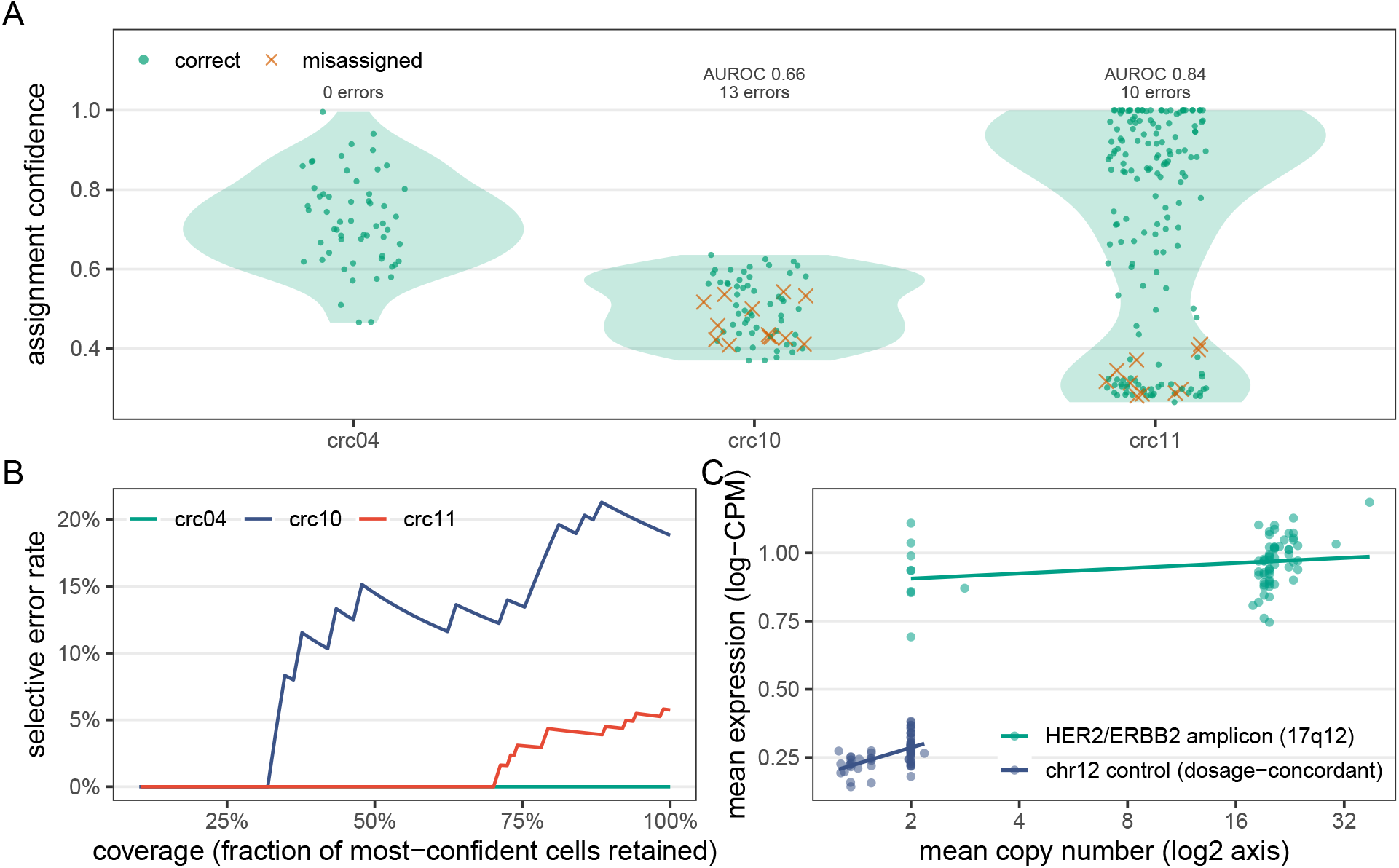
KLONE’s assignment confidence, 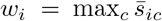, against correctness on the anchor cells under the agglomerative-raw clone definition. **(A)** Confidence of every anchor cell per patient, correct assignments as green dots with their distribution shaded and misassignments as orange crosses, with the per-patient error count and AUROC. **(B)**Risk-coverage curves, one per patient: the error rate among the most-confident fraction of cells retained. **(C)** Mean expression vs copy number for the HER2/ERBB2 amplicon and a dosage-concordant chr-12 control in `crc10` with least-squares fits.

The effect is genome-wide, not confined to one locus. `crc10`’s clones are well separated in copy number space (silhouette 0.55) but essentially unseparated in expression space (silhouette − 0.04). Global clustering therefore finds no structure in `crc10`’s transcriptome, yet KLONE still assigns its 69 anchor cells at 81.2% accuracy under the most imbalanced clone definition, well above the 65.2% majority floor: clone-specific signal survives locally even where the global structure vanishes. In short, divergent genotypes give rise to convergent transcriptomes.

These two phenomena delimit what any cross-modal method can achieve, and they account for KLONE’s residual errors. Across the same agglomerative-raw definition, KLONE misassigns 23 of the 300 anchor cells, none in `crc04`, thirteen in `crc10`, and ten in `crc11`, and the errors land exactly where the dosage signal vanishes: the `crc10` errors sit among the buffered, transcriptionally overlapping clones, and the `crc11` errors are lymph-node minority-clone cells that share partial copy number patterns with the dominant clone. We read each assignment’s confidence as its winning clone’s ensemble score, 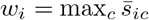 (Supp. C). Consistent with a signal limit rather than an optimization one, assignment confidence reflects this per patient (Figure 6A): in `crc11` the misassignments concentrate at low confidence (AUROC 0.84), whereas in `crc10` confidence is far less informative (AUROC 0.66) because the convergent transcriptome collapses every clone score toward the simplex floor. Read as a triage signal (Figure 6B), retaining only the most-confident assignments removes every `crc11` error but leaves `crc10`’s, whose most-confident calls still err. This points to a fundamental limit: where the RNA does not encode the genotype, no confidence score can separate right from wrong. In `crc04`, whose clones remain distinguishable in RNA, KLONE is instead perfect.

## 4 Discussion

KLONE’s lead comes from classifying each RNA cell against the labeled DNA cells rather than fitting a model that spans both modalities. The learned integrators that dominate single-cell practice (Seurat-CCA, GLUE, and MOSCOT) fit a shared structure across the modality gap without exploiting the dosage relationship and the DNA cell clone labels, so they follow the majority clone under imbalance. The two methods built for this task, TreeAlign and clonealign, do exploit dosage, but through an explicit per-gene model of expression that is not always correct. On the CRC cohort, both place every anchor cell of several runs in a single clone, resulting in catastrophic failures. KLONE instead reads the dosage signal directly off the raw similarity and departs from prior methods in three ways. It scores each RNA cell on its own similarities to the DNA cells, so the clonal distribution 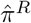 is an output rather than a constraint. Its per-clone scoring gives every clone its own neighbors, which reweighting a pooled vote cannot replicate, since a clone absent from the pooled neighborhood scores zero under any weights. Its within-*k* standardized entropy selects configurations from the unlabeled cells alone, and the resulting ensemble comes within 0.3 accuracy points of an oracle that selects with unknown RNA clone labels. Because KLONE fits no parameters and ties no assignment to the DNA clone proportions, it has nothing for a small, imbalanced, or idiosyncratically clustered dataset to destabilize. Thus, its lead widens exactly on the imbalance-aware metrics and stays flat across clone definitions.

The closed form also changes how clone assignment can be used. Since KLONE assigns a patient in 0.02 s and its weakest clone definition still exceeds every baseline’s best in accuracy, a user can run it under every candidate definition and flag the RNA cells whose assignment depends on that choice. KLONE is also not tied to copy number: it assumes only that cross-modal similarity is higher within a clone than across clones.

KLONE’s design points to two directions for future work. First, because it reads clonal identity from the *total* copy number dosage, clones distinguished only by copy-neutral or allele-specific events lie beyond its current scope, and extending the cross-modal similarity to allele-specific signals, such as B-allele frequency, is a promising route to them. Second, KLONE already emits a soft per-cell score and an entropy-based decisiveness signal. Turning these into calibrated uncertainty and abstention on the low-confidence, buffered cells it flags is a natural next step, most useful where the transcriptome’s own biology, not the assignment rule, bounds how finely genotype can be read from phenotype.

## Declaration of generative AI and AI-assisted technologies

During the preparation of this work, the authors used Anthropic Claude as supplementary tools to refine the clarity and presentation of author-written text, assist with code implementation and debugging, review mathematical formulations, and support experimental workflows. The research ideas, methodological decisions, and scientific conclusions were developed by the authors. All AI-assisted outputs were critically reviewed and verified by the authors, who take full responsibility for the final manuscript and its contents.

## Supplementary Material

Section, Figure, and Equation numbers refer to the main text. Tables and figures in this supplement are prefixed with S (e.g., Table S2, Figure S1).

## A Datasets and Preprocessing

### Patient selection

We evaluate on the colorectal cancer (CRC) cohort of Bian et al. [32], generated with single-cell triple-omics sequencing (scTrio-seq2), which jointly profiles somatic copy number, DNA methylation, and gene expression from the same cell. The original study reports paired copy-number and expression measurements for six patients (`crc01`, `crc02`, `crc04`, `crc09`, `crc10`, and `crc11`), though within each patient only a subset of cells carries both modalities. Following the benchmark of [24], we retain `crc04`, `crc10`, and `crc11`. We exclude `crc01` and `crc02` because their transcriptomes were prepared with a different library-preparation protocol than the others (the Tang protocol rather than the multiplexed scRNA-seq used for the remaining patients), which we drop to keep preprocessing uniform across patients, and `crc09` because only 13 of its cells carry both modalities, too few for a meaningful evaluation. Within each retained patient, the cells profiled in both modalities are the *anchor cells*: their shared barcodes give the ground-truth correspondence between an RNA cell and its CNV-derived clone, which we use only to score accuracy and never during assignment. Table S1 summarizes the three retained patients, ranging from the small, balanced `crc04` to the larger, extremely imbalanced `crc11`.

**Table S1:**
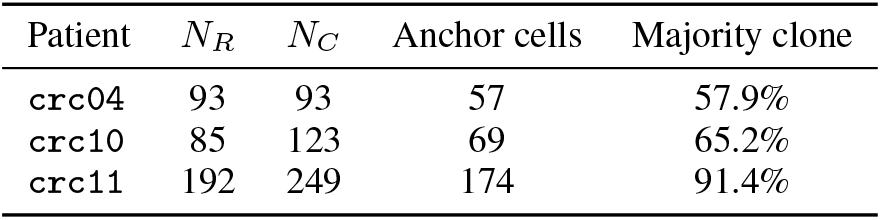
Dataset statistics for the three CRC patients. Anchor cells are those profiled in both modalities. The majority-clone column is the largest clone’s fraction of anchor cells (agglomerative-raw), from mildly skewed to extremely imbalanced.

| Patient | *N*<sub>*R*</sub> | *N*<sub>*C*</sub> | Anchor cells | Majority clone |
|---|---|---|---|---|
| crc04 | 93 | 93 | 57 | 57.9% |
| crc10 | 85 | 123 | 69 | 65.2% |
| crc11 | 192 | 249 | 174 | 91.4% |

### Preprocessing

Both modalities are restricted to the 12,894 genes measured in common, the intersection of the RNA and CNV gene sets, which is the feature space on which every cross-modal similarity is computed. Each raw cell-by-gene count matrix *X* is then processed independently per patient and per modality by the same three-step *tc-log-z* pipeline. First, total-count normalization rescales every cell to a common library size, 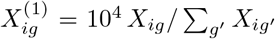, removing per-cell differences in sequencing depth. Second, a log(1 + ·) transform, 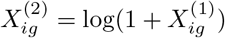, compresses the heavy right tail of the counts and copy numbers and stabilizes the variance. Third, a per-gene *z*-score across the *n* cells, 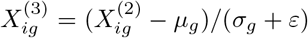 with column mean *μ*_*g*_, standard deviation *σ*_*g*_, and a small *ε* guarding the denominator, places all genes on a common scale so that no single high-dynamic-range gene dominates the similarity. Applying an identical pipeline to both modalities is deliberate. It ensures the residual cross-modal distribution gap reflects biology rather than a preprocessing mismatch, and it renders the standardized matrices 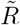 and 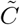 directly comparable. These outputs feed all downstream similarity computations.

### Gene selection

Each configuration operates on the top-*G* genes of a single global ranking, the HVG of Section 2.4, computed once and shared across patients. The ranking is computed on the *X*^(2)^ matrix of the preprocessing pipeline, before the per-gene *z*-score of the third step.

Write 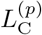 and 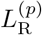 for the log(1 + ·) CNV and RNA matrices of patient *p* over the common gene set *G*. For each of the 2*P* blocks, rank genes by decreasing variance across the cells of that block, assigning rank 0 to the most variable gene. Let 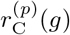 and 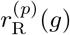 denote the resulting ranks of gene *g*. Average within each modality across the *P* patients,

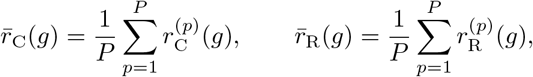

and combine the two modalities by

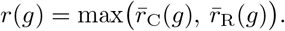

Genes are ordered by increasing *r*(*g*), with ties broken by increasing 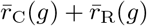 and then by gene index, and the first *G* are retained. The same gene set enters every patient’s similarity at a given *G*, so a configuration is specified entirely by the pair (*G, k*) and no gene set is tuned per patient. Two distinct centerings then act on the data, along different axes. The per-gene *z*-score above centers each gene across cells (subtracting its column mean *μ*_*g*_), whereas the Pearson correlation of Eq. (1) additionally centers each cell across the selected genes (subtracting its row mean 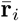 The first standardizes genes to a common scale before selection, and the second removes each cell’s mean offset at comparison time, which is what makes the similarity invariant to the per-cell shifts that differ across the two modalities.

#### Algorithm 1

KLONE

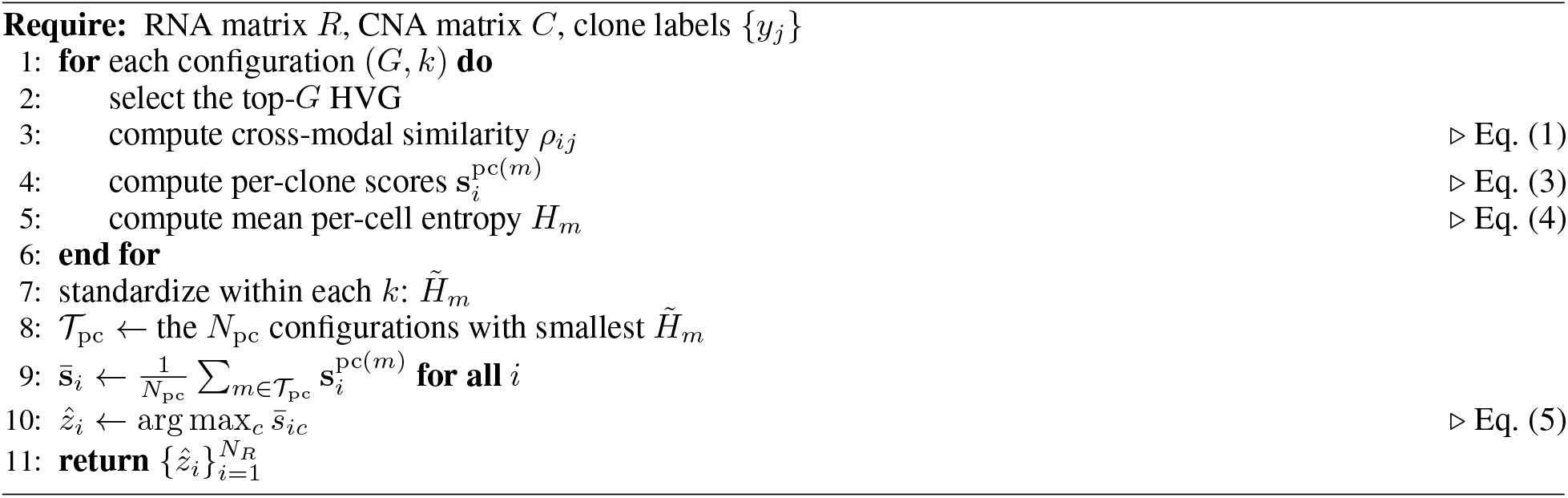

**Table S2:** KLONE hyperparameters. The method has a single free hyperparameter, *N*_pc_. Everything else is a fixed grid or a standard preprocessing choice.

| Parameter | Value |
|---|---|
| Similarity metric | Pearson correlation |
| Preprocessing | tc-log-z (total count, log1*p*, *z*-score) |
| Gene selection | HVG (RNA + CNV, all patients) |
| Per-clone gene counts *G* | {500, 750, 1000, 1500, 2000} |
| Per-clone neighbor counts *k* | {2, 3, 5} |
| Per-clone configurations *M* | 15 |
| Configurations kept *N*<sub>pc</sub> | 3 *(only free hyperparameter)* |
| Configuration selection criterion | Lowest within-*k* standardized entropy |

## B Clone Definitions

The clone labels that serve as assignment targets are obtained by clustering each patient’s scDNA-seq copy-number profiles under four definitions, formed by crossing two clustering algorithms with two copy-number transforms. *intNMF* is a model-based method built on non-negative matrix factorization that selects the number of clusters automatically, choosing the *K* that maximizes a cluster-prediction index. On the raw copy numbers, it returns *K* = 2 for all three patients, and on the log(1 + *x*)-transformed copy numbers *K* = 2 for `crc04` and *K* = 3 for `crc10` and `crc11`, a coarse major-clone resolution. *Agglomerative* clustering, a distance-based hierarchical method, does not infer *K*: we fix *K* = 4 per patient, using Euclidean distance and Ward linkage, to expose finer subclones that intNMF’s coarser partition may merge. Crossing the two algorithms with the raw and log-transformed copy numbers gives the four clone definitions (*K* = 2 to 4), and within each definition every method receives the identical labeling, so any performance difference reflects the integration method rather than the clustering.

## C KLONE Configuration

Algorithm 1 summarizes the KLONE pipeline, and Table S2 lists KLONE’s complete configuration. The method exposes a single free hyperparameter, the number of retained configurations *N*_pc_. The similarity metric, preprocessing, the (*G, k*) grid, and the label-free selection criterion are fixed across every patient and clustering, so nothing is tuned per dataset.

### Assignment confidence

For the error analysis of Section 3.8, we read each assignment’s confidence as its winning clone’s ensemble score,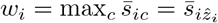: near 1 when one clone dominates the vote, near the simplex floor 1*/K* when the top clones are near-tied. It is used only to stratify assignments when analyzing errors. It never enters the assignment rule (Eq. (5)), and we set no hard confidence threshold.

## D Baseline Configurations

Each baseline is run at its recommended settings, adapted only in the label-transfer step so that clone labels flow from the DNA cells to the RNA cells rather than between two RNA assays. Preprocessing is method-specific: where a method prescribes its own normalization we follow it (MaCroDNA on raw counts, GLUE on total-count normalization and log(1+*x*), TreeAlign and clonealign on raw RNA counts and integer copy numbers, which they normalize internally), and the remaining methods (Seurat-CCA, MOSCOT) receive the same tc-log-z features as KLONE, so that any accuracy difference reflects the integration algorithm rather than the preprocessing.

KLONE, MaCroDNA, GLUE (0.3.2), MOSCOT (0.5.1), and TreeAlign (1.0) are run under Python 3.11. Seurat (5.5.1) and clonealign (1.99.3) are run under R 4.6.1.

### MaCroDNA

MaCroDNA [24] represents one-to-one matching. It computes the Pearson correlation between every RNA and DNA cell over all 12,894 common genes on the raw counts, its recommended input, and solves a maximum-weight bipartite matching that pairs each RNA cell with a distinct DNA cell so as to maximize the total correlation. When the RNA cells outnumber the DNA cells, the matching runs in successive rounds, each pairing as many cells as possible and removing them before the next, so that assignments spread across the CNV panel instead of collapsing onto a few cells. Each RNA cell then inherits the clone of the DNA cell it is matched to. The method is parameter-free, so nothing is tuned.

### Seurat-CCA

Seurat [29] represents the anchor-based label transfer that dominates single-cell multi-omic practice. We run Seurat with package defaults, except that the anchor filter is disabled since its default of 200 neighbors exceeds the number of cells in our patients. We project the two modalities into a shared space by CCA, keep the top 30 canonical components (the Seurat default), and take MNN across that space as anchors. Each RNA cell is then labeled from its anchors’ clones weighted by inverse distance, the rule used by Seurat’s `TransferData`. To isolate the algorithm from preprocessing, we give Seurat the same tc-log-z features as KLONE on the top 2,000 HVG.

### GLUE

GLUE [30] represents deep generative integration through graph-linked VAEs. We follow its recommended recipe of total-count normalization to 10^4^ and log(1 + *x*) and a Normal reconstruction likelihood suited to the resulting continuous values, and, as for the other baselines, supply the top 2,000 HVG as its highly variable features. GLUE ordinarily couples features through a prior regulatory graph, for instance, a peak-to-gene link when integrating expression with chromatin accessibility, but no such graph relates copy number to expression, so we supply the minimal self-loop graph in which each gene connects only to itself. We train for 300 epochs on CPU with a fixed seed, embed both modalities into the learned latent space, and transfer clone labels with a distance-weighted *k*-nearest-neighbor classifier (*k* = 7) fit on the CNV embeddings.

### MOSCOT

MOSCOT [31] represents OT. We cast the assignment as a cross-modality translation problem from the RNA cells (source) to the DNA cells (target) on the top 2,000 tc-log-z genes, and solve the entropically regularized transport with *α* = 0.1, entropic regularization *ε* = 0.01, and mean cost scaling. Balancedness is governed by the marginal relaxation parameters *τ*_*a*_ = *τ*_*b*_ = 1, MOSCOT’s defaults, so the coupling is balanced. Clone labels are propagated through the resulting transport plan: each RNA cell takes the clone that receives the largest transported mass, computed by multiplying the transport matrix by the one-hot CNV clone labels and taking the argmax. We also evaluated *unbalanced* OT, which relaxes the marginal constraints *τ*_*a*_ (source, RNA) and *τ*_*b*_ (target, DNA) so that the coupling need not reproduce the reference clone frequencies. To give it every chance, we grid-searched the two relaxation parameters independently, *τ*_*a*_, *τ*_*b*_ ∈ {0.1, 0.5, 0.9, 0.99, 1}, crossed with *ε* ∈ {0.005, 0.01, 0.05, 0.1, 0.5}, for 125 configurations over the twelve patient-clustering runs, counting only fully converged solves. Relaxation does not help for two reasons. First, convergence is fragile and *ε*-dependent: at MOSCOT’s recommended small regularization (*ε* ≤ 0.05) only 9 of the 25 relaxation settings converge on all twelve runs, and settings converge broadly only at *ε* = 0.5 (21 of 25), fifty times MOSCOT’s default, where the entropic coupling is heavily oversmoothed. Second, among the configurations that do converge the label-optimal one is *balanced* (*τ*_*a*_ = *τ*_*b*_ = 1, *ε* = 0.05: 80.0% accuracy, 70.3% macro-*F*_1_), barely above the default balanced OT we report in the main results (*ε* = 0.01: 78.5% accuracy, 68.5% macro-*F*_1_). No unbalanced setting improves on this balanced macro-*F*_1_: the best strictly-unbalanced converged configuration (*τ*_*a*_ = 0.99, *τ*_*b*_ = 1, *ε* = 0.5) reaches 85.4% accuracy but only 68.3% macro-*F*_1_, and it leaves the target marginal *τ*_*b*_ = 1 untouched, so it does not actually reweight the clone frequencies. Settings that do push *τ*_*b*_ down to reweight them (*τ*_*b*_ ≤ 0.9) never improve on balanced OT. Even a maximally generous oracle that uses the anchor labels to select the best converged configuration *separately for each run* reaches only 87.5% accuracy and 77.1% macro-*F*_1_, still 5.0 and 5.9 points below KLONE. Because these relaxation parameters have no label-free selection rule and their label-tuned optimum is nonetheless balanced and below KLONE, we report the balanced configuration.

### TreeAlign

TreeAlign [13] and clonealign are the two published methods built for this task. Both are probabilistic models of an RNA cell’s expression given the copy number of its clone, fit by variational inference. TreeAlign extends the clonealign model: a gene’s expected expression is its per-copy expression times the clone’s copy number, and a per-gene weight lets each gene be dosage-sensitive or dosage-insensitive. It can also use allele-specific signal and a clone phylogeny. We run it in total-copy-number mode through its clone-level interface (`CloneAlignClone`), with the DNA-defined clones as input and no phylogeny or allele-specific data, so it receives the same total copy number as the other methods. Its inputs are the raw RNA counts and the per-cell integer copy numbers over all 12,894 common genes, from which the package selects genes by its own filters. All parameters are package defaults (8 threads). TreeAlign repeats the fit 10 times with up to 400 iterations each and assigns a cell only when the same clone reaches probability 0.8 in at least 70% of the repeats (`min_clone_assign_prob` = 0.8, `min_clone_assign_freq` = 0.7). It drops clones with fewer than 10 DNA cells before fitting (`min_clone_cell_count` = 10), so no RNA cell can be assigned to them.

### clonealign

clonealign [12] models a gene’s expected expression as its per-copy expression times the copy number of the cell’s clone. We run clonealign with 8 threads at its package defaults. Its copy number input is one profile per clone, the per-gene median integer copy number over the clone’s scDNA cells. We remove genes on the X and Y chromosomes and genes with copy number 0 in any clone profile, apply `preprocess_for_clonealign` with its defaults, which also removes genes with the same copy number in every clone, and remove genes with zero expression variance after preprocessing. The fit is `run_clonealign` with its defaults and a fixed seed.

Both TreeAlign and clonealign may leave a cell unassigned when no clone is probable enough (clonealign’s default threshold is a clone probability of 0.95). We evaluate each cell’s maximum a posteriori (MAP) clone, the argmax of the fitted clone-assignment probabilities, which on our data equals each package’s call on every cell that call assigns. Section E reports the default thresholded outputs, with unassigned cells counted as wrong.

### Trivial

The trivial baseline ignores the transcriptome and assigns every RNA cell to the majority CNV clone. It is the floor that any method exploiting genuine cross-modal signal must clear, and on an imbalanced panel its accuracy already equals the majority-clone fraction.

## E Default Thresholded Outputs of TreeAlign and clonealign

TreeAlign and clonealign return a clone for a cell only when its fitted clone probability clears a threshold, and leave the cell unassigned otherwise (Section D). The main results evaluate each cell’s MAP clone. Table S3 compares it with each package’s default thresholded output, “TreeAlign (default)” and “clonealign (default)”, computed from the same fit, with unassigned anchor cells counted as wrong.

**Table S3:** Default thresholded outputs of TreeAlign and clonealign versus the MAP clones used in the main results (rows “TreeAlign” and “clonealign”). Accuracy and macro-*F*_1_ (%) are means over the three patients per clone definition, with the grand mean over the twelve runs. Unassigned anchor cells count as wrong. Coverage (%) is the fraction of the 300 anchor cells per definition that an output assigns.

|  | Method | agg-log | agg-raw | intNMF-log | intNMF-raw | Mean |
|---|---|---|---|---|---|---|
| Accuracy | TreeAlign | 63.8 | 72.2 | 56.1 | 69.3 | 65.4 |
|  | TreeAlign (default) | 63.2 | 71.2 | 41.2 | 65.0 | 60.2 |
|  | clonealign | 0.0 | 45.0 | 45.2 | 64.7 | 38.7 |
|  | clonealign (default) | 0.0 | 45.0 | 45.2 | 64.7 | 38.7 |
| Macro-*F*<sub>1</sub> | TreeAlign | 51.8 | 38.5 | 44.0 | 50.1 | 46.1 |
|  | TreeAlign (default) | 52.1 | 39.0 | 31.8 | 49.3 | 43.0 |
|  | clonealign | 0.0 | 23.1 | 35.7 | 45.4 | 26.0 |
|  | clonealign (default) | 0.0 | 23.1 | 35.7 | 45.4 | 26.0 |
| Coverage | TreeAlign (default) | 95.7 | 97.0 | 80.0 | 92.3 | 91.3 |
|  | clonealign (default) | 100.0 | 100.0 | 100.0 | 100.0 | 100.0 |

clonealign’s largest clone probability exceeds 0.99 for every cell, above its 0.95 threshold, so its default output assigns every cell and equals its MAP clone. TreeAlign’s default output leaves 105 of the 1,200 anchor-cell evaluations unassigned, 60 of them under intNMF-log, where its coverage falls to 80.0%. On every cell it does assign, it agrees with the MAP clone. Its grand-mean accuracy drops from 65.4% to 60.2% and its macro-*F*_1_ from 46.1% to 43.0%. Leaving a cell unassigned can raise a clone’s precision, so under the two agglomerative definitions the default output’s macro-*F*_1_ is slightly higher than the MAP clone’s. KLONE’s advantage remains significant against both default outputs.

Against TreeAlign (default), the McNemar split is 348*/*25 (Holm-adjusted *p* = 3.7 × 10^−73^ over the eight outputs), and the Wilcoxon adjusted *p* is at most 0.004 on both metrics. In the imbalance sweep (Section K), TreeAlign’s default output leaves 86 of the 9,390 anchor predictions unassigned and moves its mean macro-*F*_1_ by less than 1 point at every *π*_max_, and clonealign’s default output again equals its MAP clone.

## F Single-Clone Collapse of TreeAlign and clonealign

Neither TreeAlign nor clonealign clears the trivial accuracy floor in the main results. Table S4 traces this to degenerate fits. For each of the twelve patient and clone definition runs, it reports each method’s anchor accuracy next to the share of anchor cells the method assigns to its most-assigned clone. A share of 100% means the fit has collapsed and every anchor cell sits in one clone.

clonealign collapses in eight of the twelve runs. Four collapses land on a clone with no anchor cells, producing its 0% runs. Three land on a minority clone. One lands on the majority clone (`crc11` under intNMF-raw), where its output equals the trivial assignment cell for cell. A ninth run (`crc11` under agg-raw) is a near-collapse that places 97.7% of the anchors in the majority clone and scores exactly the floor. In every collapsed run, the absorbing clone is the one whose per-gene median copy number profile is closest to diploid (smallest mean |CN− 2| among the patient’s clones). This is the signature of its model design. Expression responds less than proportionally to copy number [36], so under clonealign’s strictly proportional dosage model the flattest consensus profile fits most genes best, for every cell at once. Which clone is flattest changes with the clone definition, so clonealign’s predicted clone proportions are uncorrelated with the truth (*r* = 0.01, Section L). Its non-degenerate fits are not reliable either. Only one of the four beats its patient’s floor (85.5% on `crc10` under intNMF-log). The controlled imbalance sweep shows the same behavior, with every cell sent to a flat distractor clone in each subsample (Section K).

TreeAlign collapses in five runs, the four `crc04` runs and `crc10` under intNMF-raw. The `crc10` collapse lands on the majority clone and scores the floor exactly (79.7%). The `crc04` collapses land on the DNA panel’s largest clone (41 of 93 DNA cells under agglomerative, 54 of 93 under intNMF), which holds only 24 of the 57 anchors. `crc04` is the one patient whose DNA panel and anchor set disagree on the dominant clone, so tracking the panel composition inverts the majority vote and leaves TreeAlign 15.8 points below the floor. On the other two patients, its partial minority-clone recovery lifts it above the floor (`crc10` 78.3% vs 68.8%, `crc11` 75.7% vs 70.7%). The same recovery explains the macro-*F*_1_ picture. The trivial majority vote scores 0 *F*_1_ on every minority clone, so its macro-*F*_1_ floor is only 31.6%. TreeAlign clears it (46.1%) and falls below only on `crc04` (29.6% vs 36.7%), where its collapsed assignment recovers a single clone. clonealign (26.0%) clears it only on `crc10` (43.7% vs 29.2%), the one patient where three of its four fits avoid degeneracy.

**Table S4:** Single-clone collapse. Anchor accuracy (%) per run next to the share (%) of anchor cells the method assigns to its most-assigned clone. A share of 100 marks a collapsed fit, and *†* marks a collapse onto a clone that contains no anchor cells. Floor is the trivial majority accuracy.

|  |  |  | TreeAlign |  | clonealign |  |
|---|---|---|---|---|---|---|
| Patient | Definition | Floor | Acc | Share | Acc | Share |
| crc04 | agg-log | 57.9 | 42.1 | 100 | 0.0 | 100<sup>†</sup> |
| crc04 | agg-raw | 57.9 | 42.1 | 100 | 0.0 | 100<sup>†</sup> |
| crc04 | intNMF-log | 57.9 | 42.1 | 100 | 42.1 | 100 |
| crc04 | intNMF-raw | 57.9 | 42.1 | 100 | 42.1 | 100 |
| crc10 | agg-log | 68.1 | 88.4 | 76.8 | 0.0 | 100<sup>†</sup> |
| crc10 | agg-raw | 65.2 | 79.7 | 73.9 | 43.5 | 31.9 |
| crc10 | intNMF-log | 62.3 | 65.2 | 44.9 | 85.5 | 73.9 |
| crc10 | intNMF-raw | 79.7 | 79.7 | 100 | 65.2 | 56.5 |
| crc11 | agg-log | 48.9 | 60.9 | 85.6 | 0.0 | 100<sup>†</sup> |
| crc11 | agg-raw | 91.4 | 94.8 | 89.1 | 91.4 | 97.7 |
| crc11 | intNMF-log | 55.7 | 60.9 | 93.7 | 8.0 | 100 |
| crc11 | intNMF-raw | 86.8 | 86.2 | 77.6 | 86.8 | 100 |

**Table S5:** The performance-optimal configuration is patient- and metric-dependent. Each cell gives the accuracy-optimal and macro-*F*_1_-optimal (*G, k*) as *x/y* (accuracy-optimal before the slash) over the 15-configuration grid, per patient and clustering definition. `crc04` is near-saturated (most configurations exceed 98% accuracy), so its entry is one of several near-tied optima. The optima coincide in 8 of the 12 cells and diverge only for `crc11`

| Clustering | crc04 | crc10 | crc11 |
|---|---|---|---|
| agglomerative-log | (1500, 2)/(1500, 2) | (1500, 2)/(1500, 2) | (2000, 2)/(2000, 3) |
| agglomerative-raw | (1500, 2)/(1500, 2) | (500, 5)/(500, 5) | (1000, 3)/(2000, 3) |
| intNMF-log | (1000, 2)/(1000, 2) | (2000, 2)/(2000, 2) | (1500, 2)/(2000, 5) |
| intNMF-raw | (1000, 2)/(1000, 2) | (1500, 3)/(1500, 3) | (2000, 5)/(1500, 3) |

## G Configuration Selection

Section 2.4 makes two claims about the gene budget *G* and neighbor count *k*: that no single value is best for every patient, and that neither can be tuned because the RNA cells are unlabeled. We substantiate both on the anchor cells. For each patient under each of the four clustering definitions, we score all *M* = 15 per-clone configurations (*G* ∈ {500, 750, 1000, 1500, 2000} and *k* ∈ {2, 3, 5}), recording their mean per-cell entropy, within-*k* standardized entropy, accuracy, and macro-*F*_1_.

### The optimal configuration is patient- and metric-dependent

Table S5 lists, for each patient and clustering, the two configurations that maximize anchor accuracy and macro-*F*_1_, written as *x/y* (accuracy-optimal *x*, macro-*F*_1_-optimal *y*). Three patterns stand out.

#### No single budget is optimal

The optimal gene budget spans *G* ∈ {500, 1000, 1500, 2000} and the optimal neighbor count spans *k* ∈ {2, 3, 5} : seven distinct (*G, k*) pairs appear among the twelve accuracy optima, and seven among the twelve macro-*F*_1_ optima. The field-standard *G* = 2000 maximizes accuracy in only 3 and macro-*F*_1_ in only 4 of the twelve cells. The shortfall from fixing *G* = 2000 is real but patient-specific: it costs `crc10` up to 7.0 macro-*F*_1_ points (and 3.4 to 3.8 points on two of its four clustering definitions), while the near-saturated `crc04` and the large `crc11`, whose optima already sit near *G* = 2000, lose almost nothing. A fixed default therefore does not fail loudly. It silently underperforms on whichever patient it happens to mismatch.

#### Accuracy and macro-F_1_ select different budgets under imbalance

The two optima coincide for `crc04` and `crc10` but diverge for the most imbalanced `crc11` under all four clustering definitions. The divergence is most extreme under agglomerative-raw, where two configurations reach the same 94.8% accuracy yet differ by 28 macro-*F*_1_ points (52.8% at (1000, 3) versus 81.0% at (2000, 3)). Accuracy is blind to this collapse because it is dominated by the majority clone, so a configuration chosen to maximize accuracy can quietly abandon the rare clones. This is why KLONE selects, and the paper reports, macro-*F*_1_.

*Neighborhood size tracks class balance*. The near-balanced `crc04`, saturated at most budgets, consistently prefers a small neighborhood (*k* = 2) at *G* ∈ {1000, 1500}, whereas the imbalanced `crc10` and `crc11` move to larger neighborhoods (up to *k* = 5) and shift with the clustering definition. A single fixed (*G, k*) is thus miscalibrated for at least one patient, which is exactly the sensitivity the configuration sweep is designed to absorb.

### Entropy recovers the optimum without labels

KLONE cannot read Table S5 off the unlabeled RNA cells. Instead, it averages the *N*_pc_ = 3 most decisive configurations per patient (Section 2.4), and this label-free choice recovers nearly all of the accuracy that the labeled optimum would give. Averaged over the twelve patient-clustering cells, KLONE reaches 92.5% accuracy, close to the per-patient *oracle* that selects each patient’s accuracy-optimal configuration using the ground-truth labels (92.8%), and its macro-*F*_1_ of 83.0% falls within 1.6 points of the corresponding oracle (84.6%). Two effects make the label-free selection competitive: entropy ranks the decisive, accurate configurations near the top, and averaging the three lowest-entropy ones is more robust than committing to any single configuration.

## H Ensemble Size Robustness

Section 3.6 adopts *N*_pc_ = 3 and argues that KLONE’s advantage is insensitive to it. Figure S1 and Tables S6 and S7 give the full evidence behind that claim: KLONE’s accuracy and macro-*F*_1_ on each clustering definition at every *N*_pc_ ∈ {2, 3, 5, 8, 10, 13}, with the published baselines for reference. At every *N*_pc_, both KLONE’s accuracy and its macro-*F*_1_ exceed every baseline on every clustering definition, so the lead reported in Section 3.2 does not rest on the ensemble size. The narrowest margin is macro-*F*_1_ on intNMF-log, where KLONE stays 0.4 points above MaCroDNA even at *N*_pc_ = 13, its weakest setting there. On every other definition the margin never falls below 4.1 points.

**Table S6:**
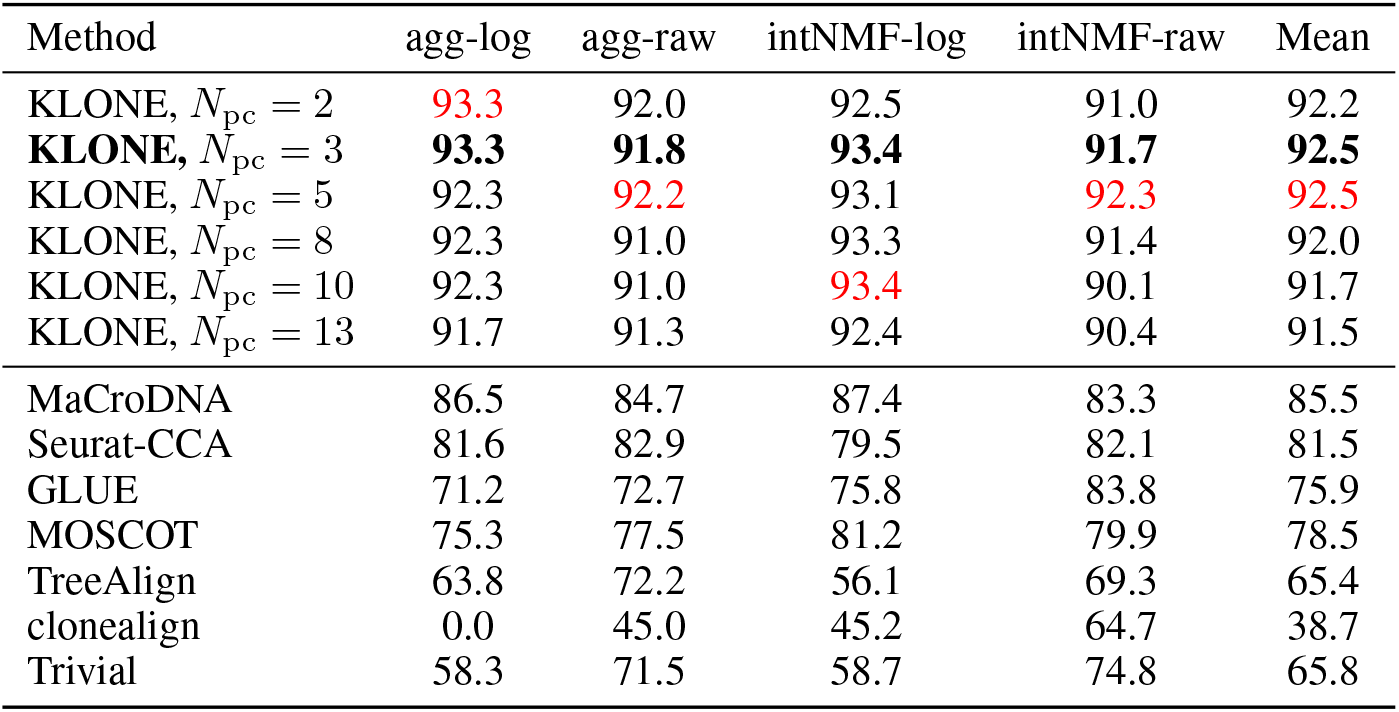
Per-clustering accuracy (%) of KLONE across the ensemble size *N*_pc_, with the published baselines for reference. At every *N*_pc_, KLONE’s accuracy exceeds every baseline on every clustering definition, so its lead does not depend on the ensemble size. The adopted *N*_pc_ = 3 is in **bold**, and the column-wise best among the remaining rows is in red.

### Accuracy is nearly constant

Table S6 shows that grand-mean accuracy varies by only 1.0 point across the entire range, from 92.5% at *N*_pc_ ∈ {3, 5} down to 91.5% at *N*_pc_ = 13. Even the lowest setting sits 6.0 points above the strongest baseline (MaCroDNA at 85.5%). Accuracy is dominated by the majority clone, whose assignment is easy and stable, so enlarging the ensemble neither helps nor hurts it appreciably. The choice of *N*_pc_ is therefore immaterial for accuracy.

### Macro-*F*_1_ is flat on the decisive plateau, then declines by design

Macro-*F*_1_ tells the more informative story (Table S7, Figure S1). For *N*_pc_ ≤ 5, it sits on a plateau, 82.6% to 83.0%, with the adopted *N*_pc_ = 3 at its top (83.0%) and its immediate neighbors *N*_pc_ = 2 and 5 within 0.4 points. Past *N*_pc_ = 5 it falls sharply, to 79.8% at *N*_pc_ = 8 and 79.0% at *N*_pc_ ∈ {10, 13}, about 4 points below the plateau. The decline is concentrated in the raw-copy-number definitions, agglomerative-raw most of all, where macro-*F*_1_ collapses from 73.9% at *N*_pc_ = 5 to 65.0% at *N*_pc_ = 8, while the log-based definitions barely move.

This decline is expected, not a fragility. KLONE averages the *N*_pc_ *most decisive* (lowest-entropy) of the 15 per-clone configurations, and only the first few are genuinely decisive. Raising *N*_pc_ past 5 forces the ensemble to average in the high-entropy, indecisive configurations, whose per-clone scores are diffuse and convolute the assignment. Accuracy absorbs the added noise because the majority clone remains an easy call, but the minority clones, which carry macro-*F*_1_, lose recall as their already-faint signal is diluted. This asymmetry is itself evidence that the entropy ranking is meaningful: the configurations it places last are exactly the ones whose inclusion degrades the metric that matters under imbalance. The region *N*_pc_ *>* 5 thus lies outside KLONE’s operating regime by construction and should not be read as a tuning frontier. Even there, KLONE’s worst macro-*F*_1_ (79.0%) still exceeds the best baseline (73.1%, MaCroDNA) by 5.9 points, so over-including indecisive configurations degrades KLONE without surrendering its lead.

## I Gene-Budget Fairness

A natural concern is that KLONE’s per-patient gene budget is an unfair advantage over baselines pinned at the field-standard *G* = 2000. It is not. Table S8 sweeps each baseline over KLONE’s grid *G* ∈ {500, …, 2000} and reports it at its own optimal budget *G*^*⋆*^. The exceptions are TreeAlign and clonealign, which take no gene panel because each selects genes from all common genes by its own filters (Section D), so we do not sweep them. MaCroDNA is included: it applies no HVG step in our main comparison, but the MaCroDNA paper [24] reports a top-2,000-gene variant, so a gene budget applies to it as well. To compare it on the same footing as the others, we sweep it over the identical grid (best *G*^*⋆*^ = 1500, 80.6%*/*67.1%), and we also list it at all common genes (85.5%*/*73.1%), its standard configuration and, unlike the HVG-native baselines, its best-performing budget. KLONE still leads every baseline on both metrics, by 7 to 13 points on accuracy and, on the imbalance-aware macro-*F*_1_, by 10 to 19 points. The gene budget is therefore not the source of KLONE’s advantage.

**Table S7:**
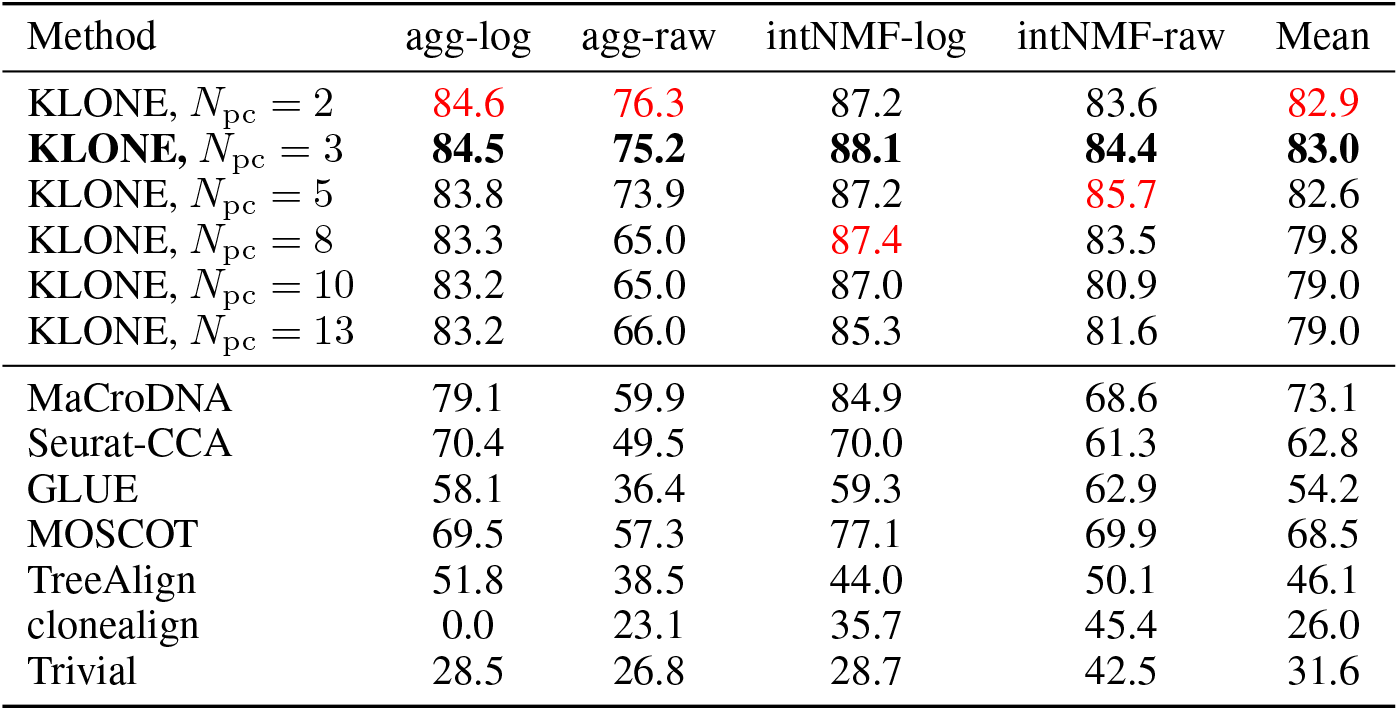
Per-clustering macro-*F*_1_ (%) of KLONE across the ensemble size *N*_pc_, with the published baselines for reference. KLONE’s grand-mean macro-*F*_1_ stays above every baseline at every *N*_pc_. The adopted *N*_pc_ = 3 is in **bold**, and the column-wise best among the remaining rows is in red.

| Method | agg-log | agg-raw | intNMF-log | intNMF-raw | Mean |
|---|---|---|---|---|---|
| KLONE, *N*<sub>pc</sub> = 2 | 84.6 | 76.3 | 87.2 | 83.6 | 82.9 |
| **KLONE,** *N*<sub>pc</sub> = 3 | **84.5** | **75.2** | **88.1** | **84.4** | **83.0** |
| KLONE, *N*<sub>pc</sub> = 5 | 83.8 | 73.9 | 87.2 | 85.7 | 82.6 |
| KLONE, *N*<sub>pc</sub> = 8 | 83.3 | 65.0 | 87.4 | 83.5 | 79.8 |
| KLONE, *N*<sub>pc</sub> = 10 | 83.2 | 65.0 | 87.0 | 80.9 | 79.0 |
| KLONE, *N*<sub>pc</sub> = 13 | 83.2 | 66.0 | 85.3 | 81.6 | 79.0 |
| MaCroDNA | 79.1 | 59.9 | 84.9 | 68.6 | 73.1 |
| Seurat-CCA | 70.4 | 49.5 | 70.0 | 61.3 | 62.8 |
| GLUE | 58.1 | 36.4 | 59.3 | 62.9 | 54.2 |
| MOSCOT | 69.5 | 57.3 | 77.1 | 69.9 | 68.5 |
| TreeAlign | 51.8 | 38.5 | 44.0 | 50.1 | 46.1 |
| clonealign | 0.0 | 23.1 | 35.7 | 45.4 | 26.0 |
| Trivial | 28.5 | 26.8 | 28.7 | 42.5 | 31.6 |

**Figure S1:**
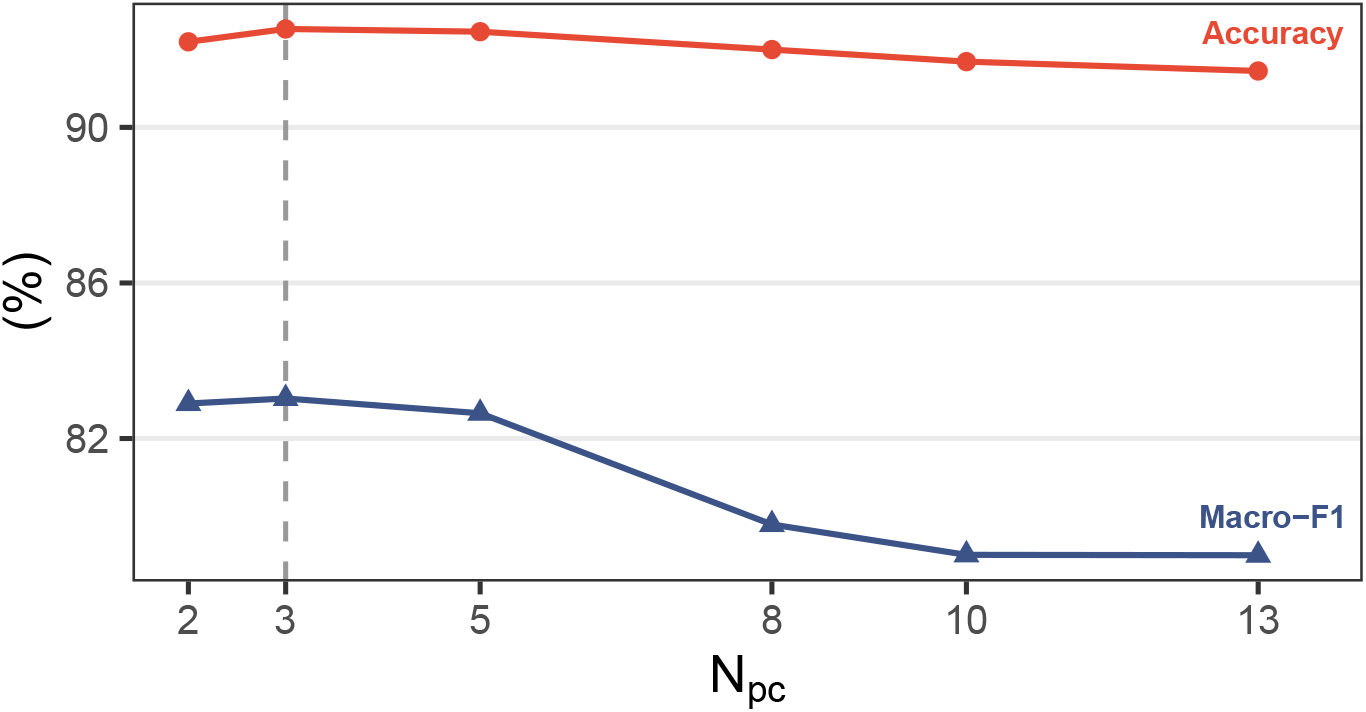
KLONE is insensitive to its lone hyperparameter *N*_pc_ within the decisive configurations: *N*_pc_ ≤ 5. As more indecisive configurations are ensembled, macro-*F*_1_ drops significantly, while accuracy stays relatively stable.

**Table S8:** Each baseline at its own optimal gene budget *G*^*⋆*^, versus KLONE, which selects *G* per patient without labels. GLUE, MOSCOT, and Seurat-CCA are swept over KLONE’s grid *G* ∈ {500, 750, 1000, 1500, 2000}. MaCroDNA appears twice: restricted to the same grid (its best is *G*^*⋆*^ = 1500), and at all common genes, its standard configuration. Grand means over twelve patient-clustering cells. KLONE leads on both metrics under either budget.

| Method | *G*<sup>⋆</sup> | Accuracy | Macro-*F*<sub>1</sub> |
|---|---|---|---|
| MaCroDNA (grid) | 1500 | 80.6 | 67.1 |
| MaCroDNA (all genes) | all | 85.5 | 73.1 |
| GLUE | 1000 | 84.6 | 66.2 |
| MOSCOT | 1500 | 79.9 | 71.5 |
| Seurat-CCA | 1500 | 82.2 | 63.6 |
| **KLONE** | per-patient | **92.5** | **83.0** |

## J Statistical Significance

We test whether KLONE’s advantage over each baseline is significant with bootstrap confidence intervals and two complementary paired tests (Table S9), using the held-out anchor clone labels for evaluation as everywhere else.

Resampling anchor cells with replacement within each run over 5,000 bootstrap replicates, the resulting 95% percentile intervals on grand-mean accuracy and macro-*F*_1_ separate KLONE from every baseline, including the strongest, MaCroDNA (accuracy [90.9, 94.0] vs [83.2, 87.7], macro-*F*_1_ [78.2, 84.8] vs [70.1, 75.3]).

**Table S9:** Paired significance tests, KLONE vs each baseline. McNemar *b/c* counts anchor cells where only KLONE(*b*) or only the baseline (*c*) is correct, pooled over the 1,200 assignments, with the two-sided binomial *p*. Wilcoxon *p* is the one-sided signed-rank value over the 12 patient ×clustering runs. Each raw *p* is followed by its Holm-adjusted value across the six baselines, separately for each test.

| Baseline | McNemar |  |  | Wilcoxon acc |  | Wilcoxon *F*<sub>1</sub> |  |
|---|---|---|---|---|---|---|---|
|  | *b*/*c* | *p* | Holm | *p* | Holm | *p* | Holm |
| MaCroDNA | 99/47 | 2.0×10<sup>−5</sup> | 2.0×10<sup>−5</sup> | 0.045 | 0.045 | 0.046 | 0.046 |
| Seurat-CCA | 134/22 | 8.7×10<sup>−21</sup> | 1.7×10<sup>−20</sup> | 0.0002 | 0.0015 | 0.0002 | 0.0015 |
| GLUE | 215/29 | 2.8×10<sup>−36</sup> | 1.1×10<sup>−35</sup> | 0.0002 | 0.0015 | 0.0002 | 0.0015 |
| MOSCOT | 162/28 | 4.0×10<sup>−24</sup> | 1.2×10<sup>−23</sup> | 0.0002 | 0.0015 | 0.0007 | 0.0015 |
| TreeAlign | 299/30 | 6.4×10<sup>−57</sup> | 3.2×10<sup>−56</sup> | 0.0024 | 0.0049 | 0.0005 | 0.0015 |
| clonealign | 634/32 | 2.8×10<sup>−146</sup> | 1.7×10<sup>−145</sup> | 0.0002 | 0.0015 | 0.0002 | 0.0015 |

### Per-cell McNemar test

Pooling all 1,200 anchor-cell assignments (the 300 anchor cells scored under the four clone definitions), we tabulate, for KLONE vs a baseline, the cells on which exactly one method is correct: *b* where only KLONE is correct and *c* where only the baseline is. Under the null of equal accuracy, the split follows Binomial(*b* + *c*, 0.5), and we report the two-sided binomial *p*-value. KLONE wins the disagreements against every baseline, all at adjusted *p <* 10^−4^ after Holm correction across the six baselines. Because the four clone definitions re-score the same 300 physical anchor cells, these 1,200 assignments are not independent, so we complement it with the coarser run-level Wilcoxon test below.

### Run-level Wilcoxon signed-rank test

Treating each of the 12 patient × clustering evaluations as one paired observation, we apply a one-sided Wilcoxon signed-rank test to the per-run accuracy and macro-*F*_1_ differences (KLONE minus baseline). Holm-correcting across the six baselines within each metric, KLONE exceeds every baseline on both accuracy and macro-*F*_1_, with adjusted *p* = 0.046 for macro-*F*_1_ and 0.045 for accuracy against the closest baseline, MaCroDNA, and *p* ≤ 0.005 against the others. The run-level margin over MaCroDNA is marginal: with only 12 correlated runs (three patients crossed with four clustering definitions), the test has little power, and KLONE wins seven to eight of them, though by larger margins than it loses. The per-cell McNemar test and the bootstrap intervals above corroborate the advantage.

## K Controlled Class-Imbalance Sweep

Section 3.4 reports a controlled experiment that raises the majority-clone proportion *π*_max_ on `crc04` while holding the patient fixed, isolating class imbalance as the single variable. Here, we detail the subsampling protocol and give the per-method numbers behind Figure 4.

### Subsampling design

Starting from the agglomerative-raw clone definition of `crc04`, we raise *π*_max_ by subsampling only the minority anchor clone. For each target *π*_max_, the majority anchor clone is kept whole and the minority anchor clone is thinned to *m* cells, which sets *π*_max_ = 33*/*(33 + *m*) on the anchor evaluation set. The CNV reference is thinned in step so that its majority proportion equals the same *π*_max_, so the imbalance a method sees in the reference is the imbalance being evaluated. Two groups are held fixed across all *π*_max_: the distractor clones, which appear in the CNV reference but carry no anchors, stay whole so the methods must still avoid mis-assigning to them, and the unlabeled RNA-only cells form a fixed-ratio random background. Sweeping *m* from 22 down to 3 yields the seven grid points *π*_max_ ∈ {0.60, 0.65, 0.70, 0.75, 0.80, 0.85, 0.92}, and we draw 30 independent replicate subsamples at each. The exact kept cells are recorded in a CSV file, so every method evaluates on identical synthetic datasets and the sampling is fully reproducible.

### Methods

At each (*π*_max_, replicate), all methods receive the same subsampled datasets and run under the default pipelines described in Sections D and C, with TreeAlign and clonealign evaluated by their MAP clones as in the main comparison. One of the 210 clonealign fits (*π*_max_ = 0.92) stopped with a non-finite ELBO inside the package’s optimizer, and we count its anchor cells as unassigned and therefore wrong. Table S10 reports mean accuracy and macro-*F*_1_ on the anchor evaluation cells over the 30 replicates. The per-cell predictions of every method on every subsample are released with the code.

**Table S10:** Controlled class-imbalance sweep on crc04 (Section 3.4). Mean accuracy and macro-*F*_1_ (%) on the anchor cells over 30 replicates, versus the majority-clone proportion *π*_max_. KLONE holds the ceiling across the range while most baselines degrade, and TreeAlign and clonealign collapse onto a single clone (see Failure modes). The best value in each column is in **bold**.

|  | π<sub>max</sub> |  |  |  |  |  |  |
|---|---|---|---|---|---|---|---|
| Method | 0.60 | 0.65 | 0.70 | 0.75 | 0.80 | 0.85 | 0.92 |
| *Accuracy* |  |  |  |  |  |  |  |
| KLONE | **99.09** | **99.74** | **100.00** | **99.92** | **100.00** | **100.00** | **100.00** |
| MaCroDNA | 74.61 | 74.05 | 73.62 | 73.56 | 72.85 | 72.91 | 73.61 |
| Seurat-CCA | 91.15 | 89.93 | 88.23 | 86.59 | 83.50 | 85.04 | 91.67 |
| GLUE | 73.39 | 84.90 | 89.43 | 80.53 | 88.54 | 91.45 | 90.37 |
| MOSCOT | 73.09 | 72.75 | 72.41 | 72.65 | 73.01 | 72.39 | 73.80 |
| TreeAlign | 40.00 | 35.29 | 29.79 | 25.00 | 80.49 | 84.62 | 91.67 |
| clonealign | 0.00 | 0.00 | 0.00 | 0.00 | 0.00 | 0.00 | 0.00 |
| *Macro-F*<sub>1</sub> |  |  |  |  |  |  |  |
| KLONE | **99.62** | **99.90** | **100.00** | **99.97** | **100.00** | **100.00** | **100.00** |
| MaCroDNA | 86.05 | 85.74 | 85.61 | 85.54 | 83.71 | 83.11 | 78.25 |
| Seurat-CCA | 90.52 | 88.24 | 83.59 | 76.55 | 57.52 | 48.52 | 48.21 |
| GLUE | 72.98 | 84.21 | 87.49 | 77.95 | 84.67 | 80.43 | 56.81 |
| MOSCOT | 78.85 | 78.24 | 77.62 | 77.02 | 75.64 | 73.61 | 61.75 |
| TreeAlign | 28.57 | 26.09 | 22.95 | 20.00 | 44.59 | 45.83 | 47.83 |
| clonealign | 0.00 | 0.00 | 0.00 | 0.00 | 0.00 | 0.00 | 0.00 |

### Failure modes

Section 3.4 reports minority-clone recall and macro-*F*_1_, and notes that the baselines’ failure modes otherwise differ. Accuracy is why they cannot be read off a single curve: over the same sweep, it stays flat for MaCroDNA and MOSCOT, dips and then rises for Seurat-CCA, moves erratically for GLUE, falls and then jumps for TreeAlign, and stays at 0 for clonealign (Table S10). Table S11 resolves this by decomposing each method’s anchor-cell predictions. Because the distractor clones carry no anchor cells, an anchor cell predicted into one of them is necessarily wrong, which separates a third error channel from the two per-class recalls.

KLONE holds 100% minority-clone recall at every *π*_max_, and its majority-clone recall stays at or above 98.5%. Its leakage into the distractor clones peaks at 0.9% in the most balanced setting and is exactly zero for every *π*_max_ *≥* 0.80. The residual error therefore moves with balance rather than with imbalance, which is the invariance claimed in Section 3.4.

MaCroDNA, MOSCOT, and Seurat-CCA fail in three distinct ways. MaCroDNA sends a constant 25 to 27% of anchor cells into distractor clones at every *π*_max_, a direct consequence of matching one-to-one against a reference that contains clones with no RNA counterpart. That constant loss pins its accuracy near 74% regardless of the clonal distribution, while its minority recall decays from 83.5% to 58.9%. MOSCOT instead leaks progressively more, from 16.4% to 24.5%, and its minority recall collapses from 72.6% to 28.9%, but its majority recall rises from 73.4% to 77.9%, and because accuracy weights the growing majority clone ever more heavily, these effects cancel and leave its accuracy flat as well. Seurat-CCA fails in the third way, by collapsing onto the majority clone. It almost never predicts a distractor clone (at most 1.5% of anchor cells) and its majority recall is at or above 97.9% throughout, but its minority recall collapses from 81.1% to 0.0%, the steepest decline of any method and a floor otherwise reached only by TreeAlign and clonealign. Because every lost minority cell lands in the majority clone, its accuracy dips only to 83.5% and then rises back to 91.7% at *π*_max_ = 0.92, where it assigns every anchor cell to the majority clone, while its macro-*F*_1_ halves from 90.5% to 48.2%. As the minority anchor clone thins, the anchors Seurat builds across the modality gap are increasingly majority-to-majority pairs, and the weighted label transfer carries the rare cells along with them.

GLUE is the exception in that its curves are erratic rather than systematic. It fits graph-linked variational autoencoders by hundreds of epochs of gradient descent on subsamples of at most 90 RNA and 75 DNA cells, far below the cohort sizes such models target, so the learned embedding is data-starved and shifts substantially from one subsample to the next. We therefore do not attribute a single mechanism to it, although its minority-clone recall does fall overall, from 57.4% to 14.4%.

**Table S11:** Failure-mode decomposition of the imbalance sweep on crc04, means over 30 replicates (%). *Distractor clones* are reference clones that contain no anchor cells, so an anchor cell predicted into one of them is always wrong. The best value in each column is in **bold** (highest recall, or fewest distractor predictions), with ties all in bold. TreeAlign’s zero distractor rate and its 100% recalls come from assigning every anchor cell to a single clone (see text).

|  | π<sub>max</sub> |  |  |  |  |  |  |
|---|---|---|---|---|---|---|---|
| Method | 0.60 | 0.65 | 0.70 | 0.75 | 0.80 | 0.85 | 0.92 |
| *Predictions into distractor clones* |  |  |  |  |  |  |  |
| KLONE | 0.91 | 0.26 | **0.00** | 0.08 | **0.00** | **0.00** | **0.00** |
| MaCroDNA | 25.15 | 25.62 | 26.24 | 26.21 | 26.91 | 26.92 | 26.11 |
| Seurat-CCA | 0.24 | 0.65 | 0.71 | 0.76 | 0.65 | 0.77 | 1.48 |
| GLUE | 11.88 | 6.27 | 2.27 | 10.38 | 7.32 | 3.16 | 4.63 |
| MOSCOT | 16.42 | 17.78 | 20.28 | 21.74 | 23.66 | 24.62 | 24.54 |
| TreeAlign | **0.00** | **0.00** | **0.00** | **0.00** | **0.00** | **0.00** | **0.00** |
| clonealign | 100.00 | 100.00 | 100.00 | 100.00 | 100.00 | 100.00 | 96.67 |
| *Majority-clone recall* |  |  |  |  |  |  |  |
| KLONE | **98.48** | **99.60** | **100.00** | 99.90 | **100.00** | **100.00** | **100.00** |
| MaCroDNA | 68.69 | 70.40 | 71.31 | 71.92 | 73.03 | 73.54 | 74.95 |
| Seurat-CCA | 97.88 | 99.49 | 99.80 | **100.00** | **100.00** | **100.00** | **100.00** |
| GLUE | 84.04 | 89.70 | 93.23 | 86.16 | 93.33 | 97.27 | 97.27 |
| MOSCOT | 73.43 | 73.94 | 74.95 | 75.96 | 77.17 | 75.96 | 77.88 |
| TreeAlign | 0.00 | 0.00 | 0.00 | 0.00 | **100.00** | **100.00** | **100.00** |
| clonealign | 0.00 | 0.00 | 0.00 | 0.00 | 0.00 | 0.00 | 0.00 |
| *Minority-clone recall* |  |  |  |  |  |  |  |
| KLONE | **100.00** | **100.00** | **100.00** | **100.00** | **100.00** | **100.00** | **100.00** |
| MaCroDNA | 83.48 | 80.74 | 79.05 | 78.48 | 72.08 | 69.44 | 58.89 |
| Seurat-CCA | 81.06 | 72.41 | 60.95 | 46.36 | 15.42 | 2.78 | 0.00 |
| GLUE | 57.42 | 76.11 | 80.48 | 63.64 | 68.75 | 59.44 | 14.44 |
| MOSCOT | 72.58 | 70.56 | 66.43 | 62.73 | 55.83 | 52.78 | 28.89 |
| TreeAlign | **100.00** | **100.00** | **100.00** | **100.00** | 0.00 | 0.00 | 0.00 |
| clonealign | 0.00 | 0.00 | 0.00 | 0.00 | 0.00 | 0.00 | 0.00 |

TreeAlign and clonealign, the two methods built for this task, fail differently again: each collapses onto a single clone. For *π*_max_ ≤ 0.75, the thinned minority clone keeps at least 10 DNA cells, and TreeAlign assigns every anchor cell to it, so its minority recall is 100%, its majority recall is 0, and its accuracy equals the minority share (40.0% at *π*_max_ = 0.60, 25.0% at 0.75). For *π*_max_ *≥* 0.80, the minority clone has only 8, 6, or 3 DNA cells, below TreeAlign’s default minimum clone size of 10 (Section D), so the package drops it before fitting and every anchor cell goes to the majority clone. TreeAlign’s macro-*F*_1_ therefore rises from 20.0% to 44.6% between *π*_max_ = 0.75 and 0.80 while it recovers no minority cell, an effect of the filter rather than robustness. The same filter drops the 4-cell distractor clone at every *π*_max_, which contributes to TreeAlign’s zero distractor rate. clonealign assigns every RNA cell of every subsample it fits to the same distractor clone, whose median copy number is 2 on every gene, so it scores 0% on every metric at every *π*_max_. Its one failed fit (*π*_max_ = 0.92) leaves 36 anchor cells unassigned, which lowers its distractor fraction there to 96.7%.

## L Clonal Distribution: Metric and Marginal Recovery

### Clonal-distribution metric

We summarize each method’s clonal-distribution recovery (Section 3.5) by a single Pearson correlation per clustering definition. For a fixed method and clustering, and for every (patient, clone) pair, we Record 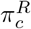, the true proportion of that patient’s anchor cells belonging to the clone, and 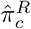, the proportion the method assigns to it (Section 2.1).

The clone set is fixed per patient by the CNV clustering, so every method is scored on the same points regardless of which clones it populates. Pooling these points across all three patients and all clones gives 12, 12, 8, and 6 points for the agglomerative-log, agglomerative-raw, intNMF-log, and intNMF-raw definitions respectively, and the correlation is taken over this pooled set. This yields one value per clustering, four per method, which Figure 5 plots with their mean and standard deviation. We pool across patients rather than correlate within each patient because a per-patient correlation would rest on only that patient’s *K* = 2 to 4 clone proportions, which is degenerate at *K* = 2 (any two points are perfectly correlated) and unstable at *K* = 3 to 4.

**Table S12:** The reference clone marginal *π*^*C*^ (over all DNA cells) versus *π*^*R*^, the clonal distribution of the evaluated anchor cells, per patient and clustering. Majority-clone proportions in %. TV is the total-variation distance (%) between the two clone distributions. The reference marginal is not the query clonal distribution.

| Clustering | Patient | maj. π<sup>C</sup> | maj. π<sup>R</sup> | TV |
|---|---|---|---|---|
| agg-log | crc04 | 44.1 | 57.9 | 20.3 |
| agg-log | crc10 | 71.5 | 68.1 | 8.3 |
| agg-log | crc11 | 41.8 | 48.9 | 11.7 |
| agg-raw | crc04 | 44.1 | 57.9 | 20.3 |
| agg-raw | crc10 | 68.3 | 65.2 | 5.0 |
| agg-raw | crc11 | 92.8 | 91.4 | 1.8 |
| intNMF-log | crc04 | 58.1 | 57.9 | 16.0 |
| intNMF-log | crc10 | 63.4 | 62.3 | 1.4 |
| intNMF-log | crc11 | 47.8 | 55.7 | 9.2 |
| intNMF-raw | crc04 | 58.1 | 57.9 | 16.0 |
| intNMF-raw | crc10 | 77.2 | 79.7 | 2.5 |
| intNMF-raw | crc11 | 87.6 | 86.8 | 0.8 |
| mean TV |  |  |  | 9.4 |

**Table S13:** The predicted clonal distribution 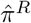 of the evaluated cells versus *π*^*R*^ and the reference marginal *π*^*C*^, grand means over the twelve patient-clustering cells (%). “pred. maj.” is the predicted proportion of the true majority clone (*π*^*R*^ 65.8, *π*^*C*^ 59.1). The TV columns are total-variation distances of 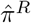 to *π*^*R*^ and to *π*^*C*^. KLONE tracks *π*^*R*^, MaCroDNA and MOSCOT track *π*^*C*^, and TreeAlign and clonealign track neither.

| Method | pred. maj. | TV to π<sup>R</sup> ↓ | TV to π<sup>C</sup> |
|---|---|---|---|
| **KLONE** | 68.6 | **3.5** | 11.1 |
| MaCroDNA | 62.0 | 11.1 | 7.8 |
| MOSCOT | 57.1 | 11.3 | 4.8 |
| Seurat-CCA | 70.3 | 14.6 | 13.5 |
| GLUE | 73.5 | 16.7 | 19.3 |
| TreeAlign | 52.5 | 32.7 | 31.8 |
| clonealign | 29.0 | 58.9 | 51.8 |

Section 2.1 argues that a method should assign each cell from its own cross-modal evidence, not from the reference panel’s clone frequencies *π*^*C*^. The rest of this section substantiates the two quantitative claims behind that argument: that *π*^*C*^ differs measurably from the clonal distribution *π*^*R*^ of the evaluated cells, and that KLONE’s 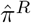 tracks *π*^*R*^, while the frequency-constrained baselines track *π*^*C*^.

### The reference marginal is not the query clonal distribution

The datasets are only partially paired: the anchor cells carry both modalities, but each patient also contains cells profiled in a single modality (36 to 75 DNA-only cells here), and the DNA reference and RNA query are otherwise separately sampled. The reference frequency 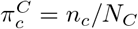 (over all DNA cells, the quantity that matching and transport import) therefore need not equal *π*^*R*^, the clonal distribution of the evaluated anchor cells. Table S12 shows it does not: the majority-clone proportions differ by up to 13.8 points, and the two distributions differ by a mean total-variation distance of 9.4% (up to 20.3%). A frequency-constrained method thus imports a frequency that is measurably not the query’s.

### KLONE recovers *π*^*R*^, frequency-constrained baselines recover *π*^*C*^

For each method, we compute 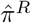 over the evaluated cells and compare it, by total-variation distance, to *π*^*R*^ and to *π*^*C*^ (Table S13, grand means over the twelve patient-clustering cells). KLONE lands within 3.5% TV of *π*^*R*^, at least three times closer than any baseline, because its per-clone scoring reads the proportions off the per-cell evidence rather than a prior. Balanced OT (MOSCOT) and one-to-one matching (MaCroDNA) instead sit closer to *π*^*C*^ than to *π*^*R*^ (TV to *π*^*C*^ below TV to *π*^*R*^), the signature of constraining the induced marginal to the reference panel, while Seurat-CCA and GLUE overshoot past both toward the majority extreme. TreeAlign and clonealign land far from both (TV above 30%) and underweight the true majority clone, because in several runs they place every anchor cell in a single clone other than the majority (TreeAlign under all four definitions of `crc04`, clonealign for all three patients under agglomerative-log). Because the true majority reaches 91% on `crc11` yet is recovered by a method that imports no clone prior, the proportions come from the likelihood, not from a prior.

